# CUGIC: The Consolidated Urban Green Infrastructure Classification for assessing ecosystem services and biodiversity

**DOI:** 10.1101/2022.05.16.492061

**Authors:** Joeri Morpurgo, Roy P. Remme, Peter M. Van Bodegom

## Abstract

Green infrastructure (GI) classifications are widely applied to predict and assess its suitability for urban biodiversity and ecosystem service (ES) provisioning. However, there is no consolidated classification, which hampers elucidating synthesis and consolidated relationships across ES and biodiversity.

In this research, we aim to bridge the gap between urban GI research on ES and biodiversity by providing a standardized common classification that enables consistent spatial analysis.

We analyzed GI classifications used across five ES and four taxa in scientific literature. GI classes were analyzed based on name, definition and characteristics. Results were used to create a novel classification scheme accounting for both ES and biodiversity.

We show that many GI classes are unique to a ES or taxon, indicating a lack of multifunctionality of the classification applied. Among the universally used classes, diversity in their definitions is large, reducing our mechanistic understanding of multifunctionality in GI. Finally, we show that most GI classes are solely based on land-use or land-cover, lacking in-depth detail on vegetation. Through standardization and incorporation of key characteristics, we created a consolidated classification. This classification is fully available through openly-accessible databases.

Our consolidated standardized classification accommodates interdisciplinary research on ES and biodiversity and allows elucidating urban biodiversity and ES relationships into greater detail, facilitating cross-comparisons and integrated assessments. This will provide a foundation for future research efforts into GI multi-functionality and urban greening policies.

## 1. Introduction

Green infrastructure (GI) is essential for biodiversity and the provisioning of ecosystem services (ES) in the urban environment (Escobedo et al., 2019; Sun et al., 2020), where ES are contributions of ecosystems to human wellbeing (TEEB, 2010). GI forms the basis for nature-based solutions to major urban challenges such as climate change, biodiversity and human wellbeing (Escobedo et al., 2019; IUCN, n.d.). Policy makers are increasingly targeting GI as a key method to tackle such challenges simultaneously and uniformly (Liberalesso et al., 2020). For example, the European Commission defines and uses GI as: “a strategically planned network of high quality natural and semi-natural areas with other environmental features, which is designed and managed to deliver a wide range of ecosystem services and protect biodiversity in both rural and urban settings” (European Commission, 2013). Unfortunately, while a multi-functionality from GI is implied by this definition, in practice the complex relationships between these ES and biodiversity and their relation to GI are hardly accounted for.

Both biodiversity and ES are considered important to protect and enhance, both for human wellbeing and for the health of our planet (Brondizio et al., 2019; Chaplin-Kramer et al., 2019). Urban GI plays an important role in supporting them. GI supports ecosystem functioning and its biota through supporting, increasing or conserving key species and taxa (Dearborn & Kark, 2010; McKinney, 2002; Niemelä, 1999). Additionally, GI provides important urban ES, such as heat reduction, air purification, water regulation, and supports both physical and mental wellbeing (Luedertiz et al., 2015). Such urban ES are well studied and associated with the increase in human wellbeing or a reduction of economic damages (Bolund & Hunhammar, 1999; Luedertiz et al., 2015; Meerow & Newell, 2019). While GI presumably contributes to both biodiversity and ES and that nature – and thus GI - should be beneficial for both humans and biodiversity (Heymans et al., 2019), they are commonly investigated separately, in different studies. Unfortunately, the divide between urban biodiversity and ES studies also translates into the usage of dissimilar GI classifications and definitions.

Existing classifications aim to provide insight into the spatial patterns of GI in relation to its respective services or taxon, but are often created ad hoc and do not relate to other ES or taxa. This creates a large diversity of incomparable definitions that are being used to classify GI. For example, the GI class grassland has been classified by (i) their dominant species (Miralless-Guasch et al., 2019), (ii) height of vegetation (Ng et al., 2012), (iii) land cover maps (Tiwari and Kumar, 2020), or (iv) simply based on expert opinion (Morelli et al., 2018). Classifying GI with a breadth of different definitions results in poor cross-comparability, impeding elucidation of driving factors in complex relationships (Bartesaghi Koc et al., 2017; Chatzimentor et al., 2020; Wang & Banzhaf, 2018).

From a scientific perspective, these inconsistencies seriously impede our understanding of the relations, synergies, and potential trade-offs, among biodiversity and ES in the urban environment (Fineschi and Loreto, 2020; Manning et al., 2018; Schwarz et al., 2017). This lack of interdisciplinary knowledge transaction limits the efficiency of multifunctional urban environmental planning (Daily et al., 2009). Additionally, the lack of consistency between GI classifications hinders integral policy, planning, and monitoring by authorities, as they create ambiguity (Garmendia et al., 2018). Therefore, it is paramount to researchers, policy makers, and urban planners to have a comprehensive and consistent understanding of GI to easily and effectively communicate new findings.

To better understand the complex GI-ES-biodiversity relationship we need mechanism-relevant standardized urban GI classifications and definitions (Bartesaghi-koc et al., 2017; Matsler et al., 2021 Schwarz et al., 2017). Creating such a harmonized classification is a challenging process as the relevant mechanisms, spatial and temporal scale that drive ES or biodiversity are vastly different, and need to be accounted for. Nonetheless, notable efforts to standardize and consolidate GI classifications have been made. For instance, the Green Infrastructure Typology (Bartesaghi-koc et al., 2017), the Urban Vegetation Structure Types (Lehmann et al., 2014), and the High Ecological Resolution Classification for Urban Landscapes and Environmental Systems (Cadenasso et al., 2007) have all been proposed as a general GI typology. Unfortunately, these classifications either solely focus on ES, are laborious to create, or contain ambiguous definitions (Appendix 1). Among the existing GI typologies, little attention has been given to the ecological mechanisms at play in GI that provide ES or support biodiversity (Young et al., 2014). This is further exacerbated by the lack of a combined typology for both biodiversity and ES, hindering research and policy that aim to incorporate them simultaneously. A key area of interest that would benefit from a combined typology relates to synergies and trade-offs between urban ES and biodiversity, that remain largely unexplored to date (Schwarz et al., 2017). To that end, and as a solid foundation for future integrated research, a consistent and broad classification system for urban GI is needed.

In this research, we aim to bridge the gap between urban ES and biodiversity research by creating a harmonized GI classification based on previous ES and biodiversity research. This allows future research to elucidate their relationship, and policy makers and practitioners to plan and manage GI more holistically. We conducted a semi-systematic literature review focused on GI classifications at three levels: (i) names, (ii) definitions, and (iii) characteristics. For this review, we selected four ES that play central roles in urban resilience and sustainability and are commonly studied: water regulation, heat reduction, air purification, mental and physical wellbeing (Keeler et al., 2019; Schwarz et al., 2017; Veerkamp et al., 2021). We also study four taxa that together reflect broader urban biodiversity: mammals, plants, birds, arthropods (Beninde et al., 2015; Chatzimentor et al., 2020; Schwarz et al., 2017).

Based on an in-depth multi-level overview on the GI classifications used in previous studies, we constructed a new evidence-based consolidated GI classification that unifies urban ES and biodiversity. Importantly, this new classification is applicable with remote-sensing data to accommodate analysis in cities across the world. The resulting classification aims to accommodate future multifunctional research concerning ES and biodiversity in the urban environment. Moreover, by providing definitions for GI types and linking them to ES and biodiversity, this classification allows for cross comparisons and can guide policymakers aiming to optimize of GI in the increasingly dense urban environment (Matsler et al., 2021).

## 2. Methods

### 2.1 Data collection

To identify and analyze current usage of GI typology, we reviewed the literature on GI in relation to ES and biodiversity, published between 2011-2021. We chose this time-frame as the number of GI-related publications greatly increased after 2010, as well as to include more recent fine scale data from Sentinel-2 and OpenStreetMap for analyzing urban contexts (Ludwig et al., 2021; Matsler et al., 2021). We applied a semi-systematic review by using thematic saturation on the Web of Science (Guest et al., 2020; Gusenbauer & Haddaway, 2020; Appendix 2). We chose this method over other review methods, such as PRISMA, as our goal is extract qualitative data and consolidated it into a new classification. In particular, the chosen saturation method allows researchers to rapidly identify themes in the literature, while also enabling to numerically quantify the confidence in their results.

We used one search query per service or taxon applying it to the title, abstract and keywords on the Web of Science (Appendix 3). We chose to rank papers by the number of citations to prioritize the most representative GI research, yet we acknowledge that they are likely older and biased to coarse resolutions and research in the northern hemisphere (Matsler et al., 2021). We included papers that: I) contained two or more GI classes, to exclude studies focusing solely on one vegetation indicator, II) focused on a single ES or taxon, to bring out GI classes that uniquely link to a single ES or taxon, and III) were focused on urban land in the studied area (see Appendix 2 for more details).

In the initial search we found 3990 articles, of which we analyzed 143 based on the previously listed inclusion criteria. 75 studies focused on ES and 68 on biodiversity (see Appendix 4 for the list of analyzed articles). After collection, we analyzed all GI classes (n = 564) for each ES and taxon for three aspects (Fig. 1; elaborated in sections 2.3-2.5): First, we evaluated the most commonly used class names. Second, we analyzed the diversity and overarching themes in GI definitions. Third, we assessed GI class characteristics. Based on these analyses, we created a harmonized classification of GI that is applicable to both biodiversity and ES assessments (section 2.6).

**Figure 1.**
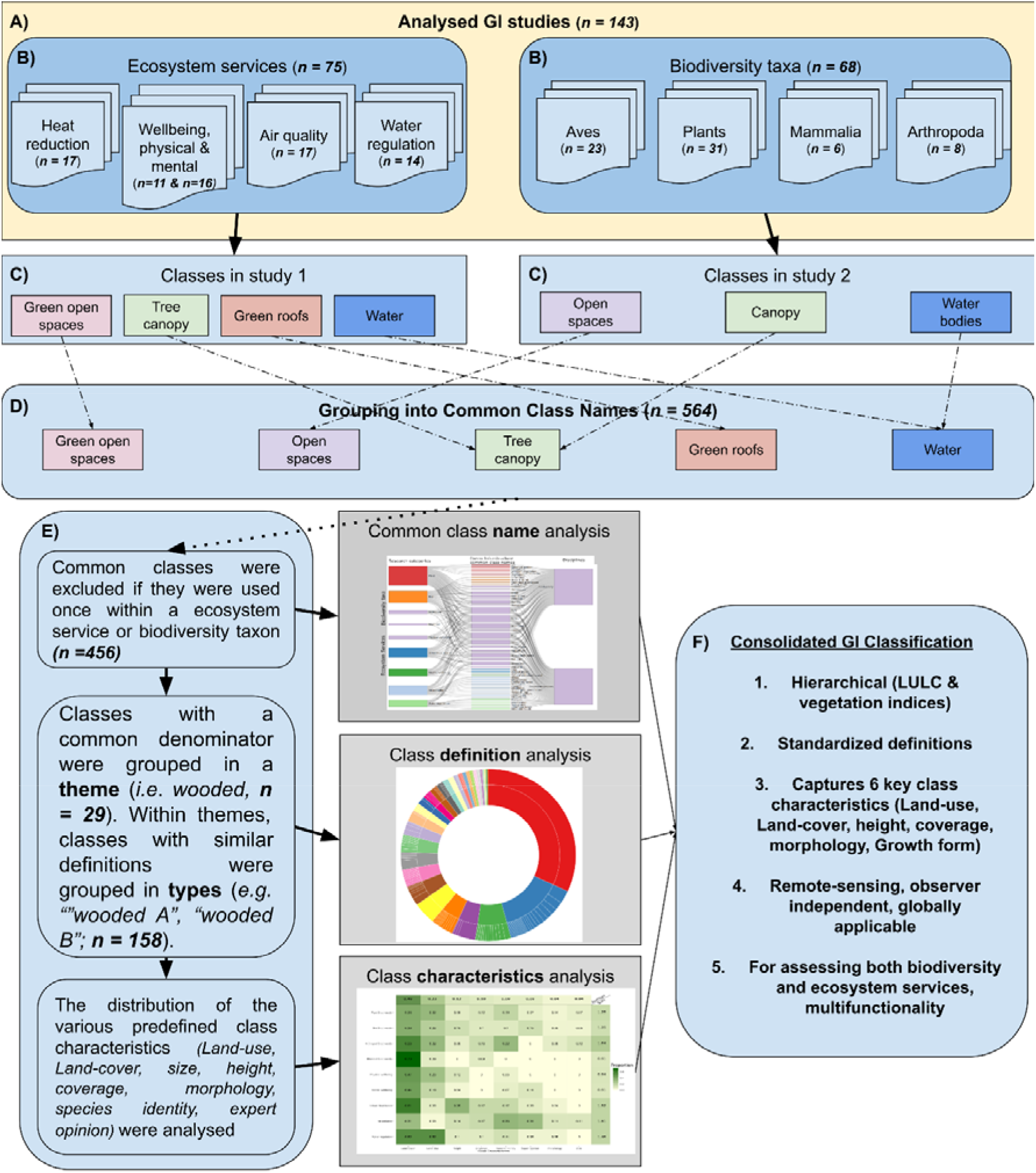
Flowchart of data collection and analysis. The top shows the number of included studies (A) and number of included studies across ES and biodiversity taxa (B). For each study, the classes identified were listed (C) illustrated with two arbitrary examples and common class names were identified (D). A three tiered analysis (E), involving class names, definition, and characteristics followed to create an evidence basis for the consolidated GI classification (F).

### 2.2 Saturation

After collecting eight papers for a service or taxon, we analyzed its thematic saturation for GI classes (Guest et al., 2020). In short, thematic saturation was measured by: I) base size, II) run length and III) New Information Threshold (NIR, methods in Appendix 2). The NIR index was used to indicate how likely it is to find new information with additional sampling. A high percentage indicates that the next sample is likely to contain a lot of new information. We used a base size of five papers, run length of three papers, and a NIR of 10%. This method is mainly applied for interviews, but we deem this approach useful as there is no agreed upon method to measure saturation for our kind of data (Saunders et al., 2018). Grey and blue infrastructure classes were analyzed but are beyond the scope of this research and therefore put in the appendices (Appendix 5).

Thematic saturation was reached from literature review on ES; urban heat island (NIR = 0; n = 17), water regulation (NIR = 0.09, n =14), and air pollution (NIR = 0.10, n = 17), and for the arthropod taxon (NIR = 0.10, n = 8). In contrast, saturation was not reached for the bird taxon (NIR = 0.24, n = 23), the plant taxon (NIR = 0.26, n = 31), the mammal taxon (NIR = incalculable, n = 6), mental health (NIR = 0.33, n = 16) and physical health ES (NIR = 0.50, n = 11) when the literature from our search results had been fully exhausted, thus providing the most up-to-date results possible.

### 2.3 Class name analysis

In order to create an overview of GI classes that are representative for a service or taxon, we used a Sankey diagram to link commonly used GI class names to ES and taxa. We decided to discard class names that were only used once for a service or taxon to only include classes that are well-linked with an ES or taxon. After exclusion of the single-use classes, we calculated the frequency of GI class usage for ES or biodiversity. In particular, this method allows us to infer which classes are universally used and which are uniquely used for a single service or taxon.

### 2.4 Class definition analysis

To more comprehensively investigate the diversity of urban GI classifications, we also analyzed the class definitions used. For each class, a definition was collected from its respective paper. Definitions were analyzed in a two tiered process which consisted of the creation of definition themes and types. The first tier, themes, consisted of GI classes of which definitions could be largely grouped by a common denominator. For example, the theme “Grassland” contained all classes with a common denominator, being: “green spaces with grass as dominant vegetation”.

The second tier, named types, are nested within a theme. For example, within the “Grassland” theme, consisting of 30 classes, we identified multiple types of common definitions (n = 15). Where all classes in type A (n = 5) define Grassland as “Area covered mostly by grass or lawn”, classes in type B (n = 3) define Grassland as “grassland” using land cover maps. While for example classes in type E (n = 2) define Grassland as “Grass average 50cm height”. Following this procedure, we argue that types indicate the variety in operationalization of GI classes within a theme.

### 2.5 Class characteristics analysis

In order to more thoroughly understand how the included classifications were designed, we decided to score the definitions of individual classes, from every classification, included in the analysis based on eight characteristics. These eight key class characteristics were identified based on commonly used features in GI classifications. These include: (1) land use, (2) land cover, (3) size, (4) height, (5) coverage, (6) morphology, (7) species identity, and (8) use of expert opinion (See appendix 6 for more details). We applied a binary score per characteristic for every GI class their definition. (1= present, 0=absent). For example, a grassland class defined as “vegetation below 0.50m and larger than 1 ha” would score 1 for height and size, but 0 for the other characteristics.

Based on the binary scores, we calculated the average presence for each characteristic among classes for each service or taxon. As such, a value of 1 would indicate that every class within a service or taxon explicitly mentions that characteristic, while a value of 0 indicates none of the classes mention a characteristic explicitly. Second, we compared the total usage of explicit class characteristics among ES and taxa. We summed the values from all characteristics by service or taxon, indicating how characteristics-dense classes are on average within a service or taxon. Values of eight would indicate that every class within a taxon or service mentions all eight characteristic explicitly, while values of 0 would indicate none of the classes mention any class characteristic explicitly. The results were visualized by a heat map showing the difference in the usage of explicit class characteristics by service or taxon.

### 2.6 Consolidated Urban Green Infrastructure Classification (CUGIC)

To create a consolidated urban GI classification, which we termed CUGIC, we started with the identified themes (n = 29) from the class definition analysis (section 2.4) as the basis for the consolidated classification (e.g., “Green roof” being: “*Vegetation that partially or fully covers a roof*”). These themes reflect key aspects of GI identified in literature. In parallel, from the analysis of class characteristics (section 2.5), we chose to include *LU* & *LC* classes as they were the most abundant, and chose to include the characteristics that cover structure of vegetation (*height, coverage, morphology, and Growth form*). While vegetation structure was not used as consistently as LULC across ES and taxa, they play a key role in describing vegetation variation within LULC classes. We excluded the two final characteristics, *expert opinion* and *size*, as we prioritized observer-independent sampling (Simensen et al., 2018), and the *size* characteristic was used both infrequently and inconsistently across definitions.

Next, we aimed to translate the GI themes into classes that could be easily quantified with globally freely available data sources, and that allowed for links to the six chosen characteristics (see appendix 6 for details). These six characteristics were grouped into two separate data layers. One data layer covered LULC (land-use, land-cover) and the other covered vegetation structure (height, coverage, morphology, and species identity (simplified to Growth form)). The combination of both generally increases model performance (Shen et al., 2021; Chaves et al., 2020). Moreover, this combination also ensures that the classification was applicable with both remote-sensing (e.g., LiDaR, Normalised Difference Vegetation Index (NDVI), etc.) and LULC data.

The LULC data layer originally contained 22 themes and was reduced to 16 LULC classes in the CUGIC. We chose to exclude two themes. One excluded theme described a specific virtual reality GI courtyard set-up, and was excluded on the basis that it was only used in a simulation study (Huang et al., 2020.). The other theme described green open spaces. However, this theme was nearly all-encompassing, and was excluded on the basis that it was not mutually exclusive with the other themes. Further, we made three consolidations covering 7 themes. First, three themes (remnant, spontaneous, and lot) which all classified vegetation that grew without management were combined into one class: *remnant vegetation*. Second, we combined raingarden and bio retention basin into one class: *raingarden* as they are synonymous. Third, we combined residential green and garden into one class: *residential green* & *gardens*, as they both comprise of private green space and given that flowers, the focus of the garden theme, are not easily distinguished from residential green by remote-sensing.

The vegetation structure section contained 7 themes and were increased to 41 vegetation classes in the CUGIC. The themes covering vegetation mainly comprised of structural thresholds (coverage or height), morphology (number of layers), and growth form (grass, shrub, trees, evergreen or deciduous). We chose to create a novel set of classes based on these themes. Leaf habit information (i.e., (ever)greenness and deciduousness) was included as it is often found in LULC maps. The CUGIC therefore defines vegetation classes by I) height, II) coverage density, III) whether it has one or more layers, and IV) if it contains evergreen or deciduous trees.

## 3. Results

### 3.1 Class name analysis

Analyzing common class names that were present twice or more in a specific service or taxon resulted in a list of 49 unique common class names (Fig. 2). Class names focusing on large commonly urbanized vegetation were most frequently used, being forest (n = 35), park (n = 32) and tree (n = 31). GI classifications for plants have several classes that are unique to its topic (Fig. 2, red boxes). These mostly include classes that concern land management (remnant, spontaneous, plantations). Also water regulation and air pollution have several unique classes (Fig. 2, light blue and light green boxes). Air pollution has unique classes focusing on vegetation morphology (one layer, multiple layers, hedge), and on growth form (broad leaved, coniferous, deciduous). Water regulation shows, among all connections, five unique classes that are strongly related to constructed land-use (bioswale, bio retention, rain garden, infiltration basin, detention basin). These results indicate that there is a large diversity of class usage amongst services and taxa with several GI classes being completely unique to a service or taxon, while only few are used almost universally.

**Figure 2.**
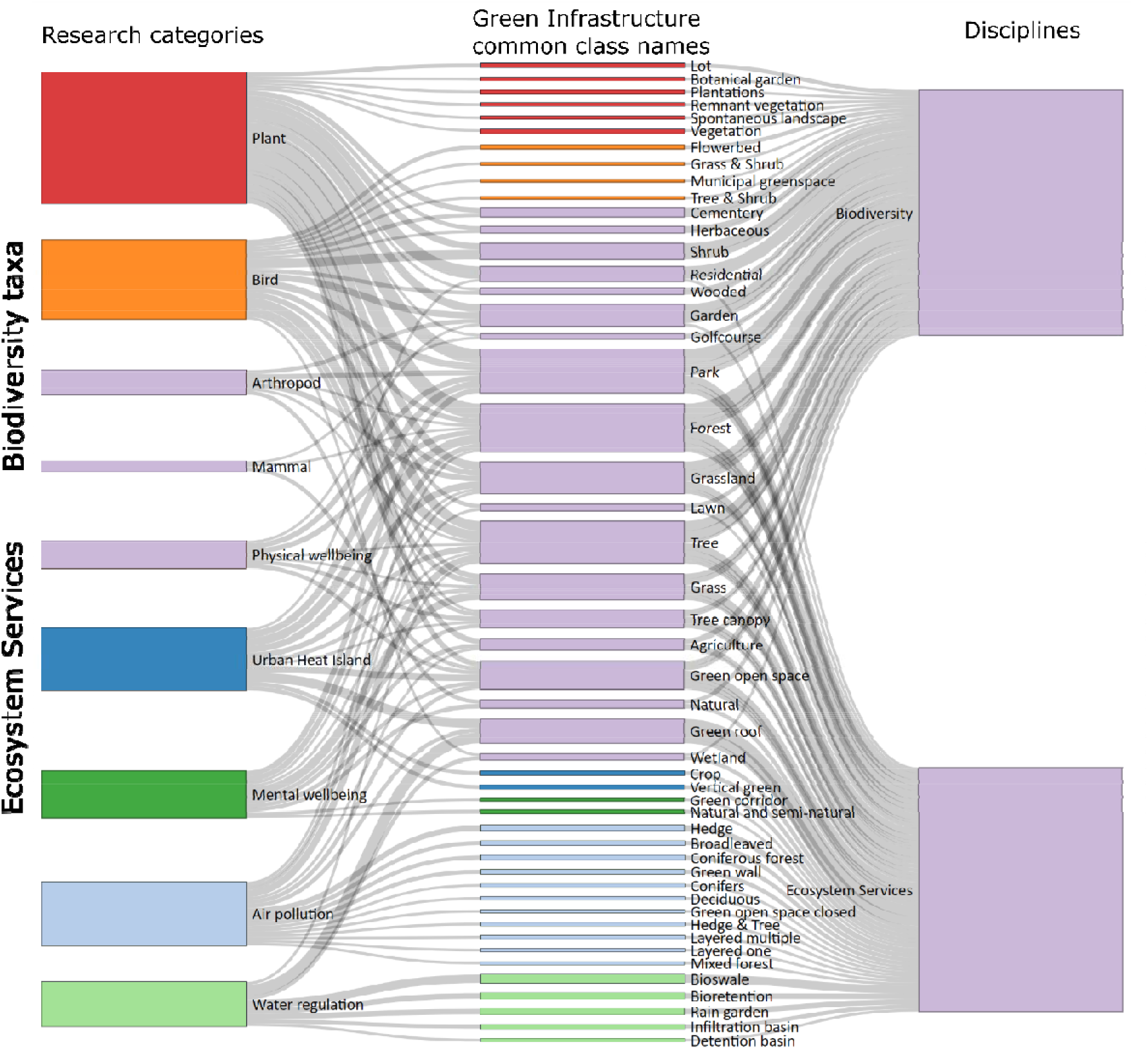
Sankey diagram showing the links between services or taxon and GI class names. Nodes in the middle are ordered with the least frequently used GI classes on the edges, while most frequently used GI classes are in the center. The thickness of the nodes and flows is based on frequency. The colors of the nodes represent either a unique or multiple connection, with the purple color representing multiple connections across biodiversity and ES assessments. Other colors represent an unique connection between a taxon or ES and a GI class (e.g. Urban heat island being uniquely connected to crop and vertical green).

### 3.2 Class definition analysis

The definition of each GI class (n = 456) was analyzed. A large proportion of the classes were left undefined in their respective research paper (n = 145, 31.9%), meaning that a clear class identification and/or description were missing. A smaller portion was unavailable, e.g., behind a paywall, a dead link, or in a different language than English (n = 19, 4.2%). From the defined classes, we identified twenty-nine themes which contained 158 different types (Fig. 3, appendix 7). This indicates that a large diversity of GI definitions is used within one theme.

**Figure 3.**
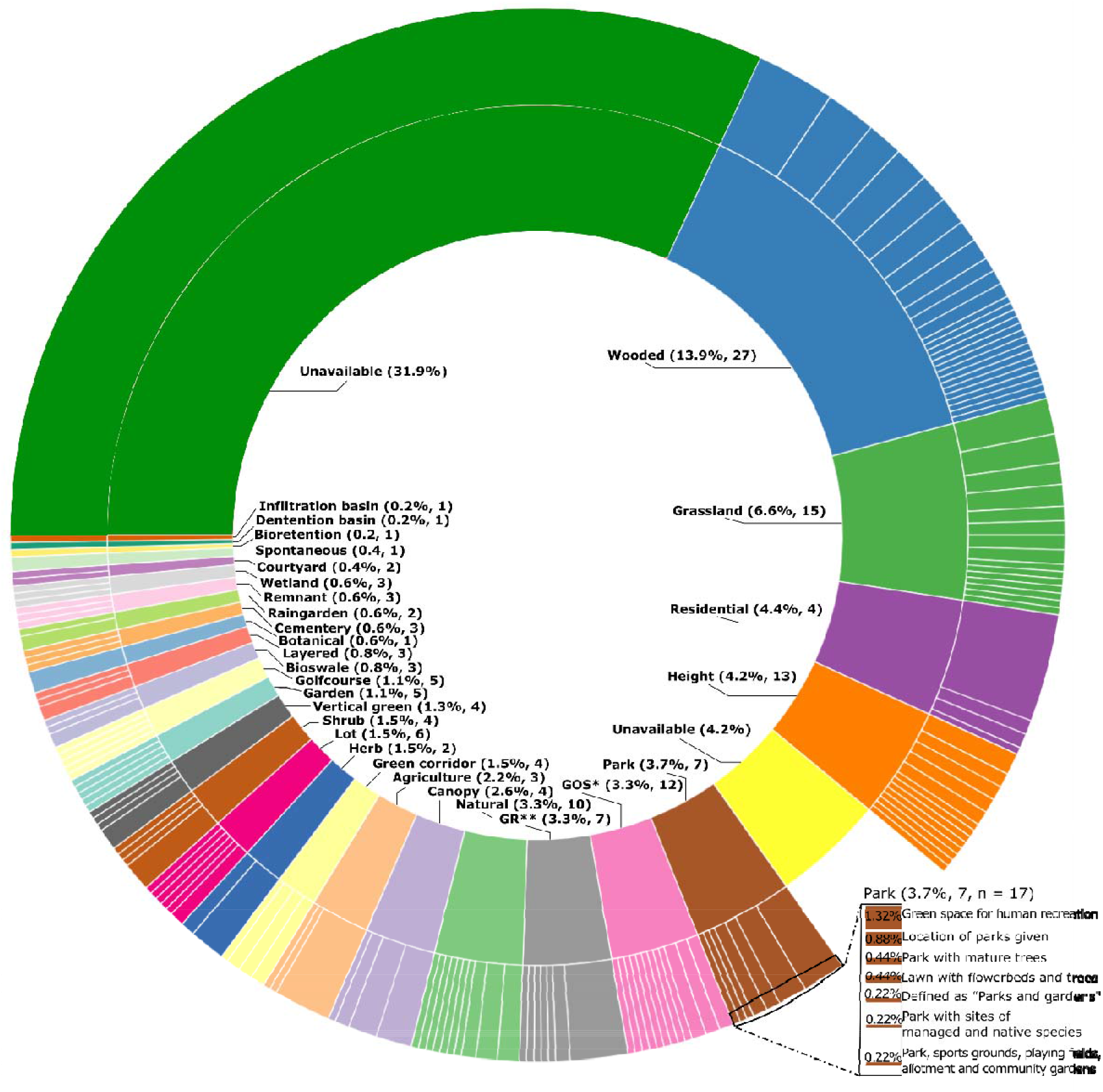
Sunburst diagram of the GI definition theme and types. The inner layer contain themes, and the outer layer types, portraying the number of GI types in one theme. Zoomed in is the Park theme, showing the seven definition types within. * - Green Open Space. ** - Green Roof.

The largest identified theme was “wooded” which contained 27 types that all differently defined land with substantial or continuous cover by woody vegetation, and could be a mix of deciduous, coniferous or evergreen species. The smallest themes were “infiltration basin” and “detention basin” with both only one type, which was based on expert opinion. Two themes were identified for using solely height or vegetation coverage thresholds. These themes “height” and “canopy coverage” were used as indicators for vegetation, yet the thresholds varied greatly between studies (e.g. forest being either >5m, >10m, >15m). These results indicate that regularly measured GIs contain a broad variety of definitions.

### 3.3 Class characteristics analysis

We found that the characteristics most GI classes used, across services and taxa, are land-use (49%) and land cover (23%), while the characteristics morphology (4%) and size (4%) are used least (Fig. 4). The heat map illustrates that there is more difference among characteristics than among services or taxa (Fig. 4). This indicates that GI classes among ES and taxa have similar characteristics (despite some unique class names and dissimilar definitions). Among the ES and taxa, we find that mental health (0.91), physical health (0.94), and mammal (0.91) studies use on average the least number of characteristics per class, while arthropod research (1.64) defines on average the most characteristics per class. This indicates that most ES or taxa studies consider only one characteristic of GI classes explicitly.

**Figure 4.**
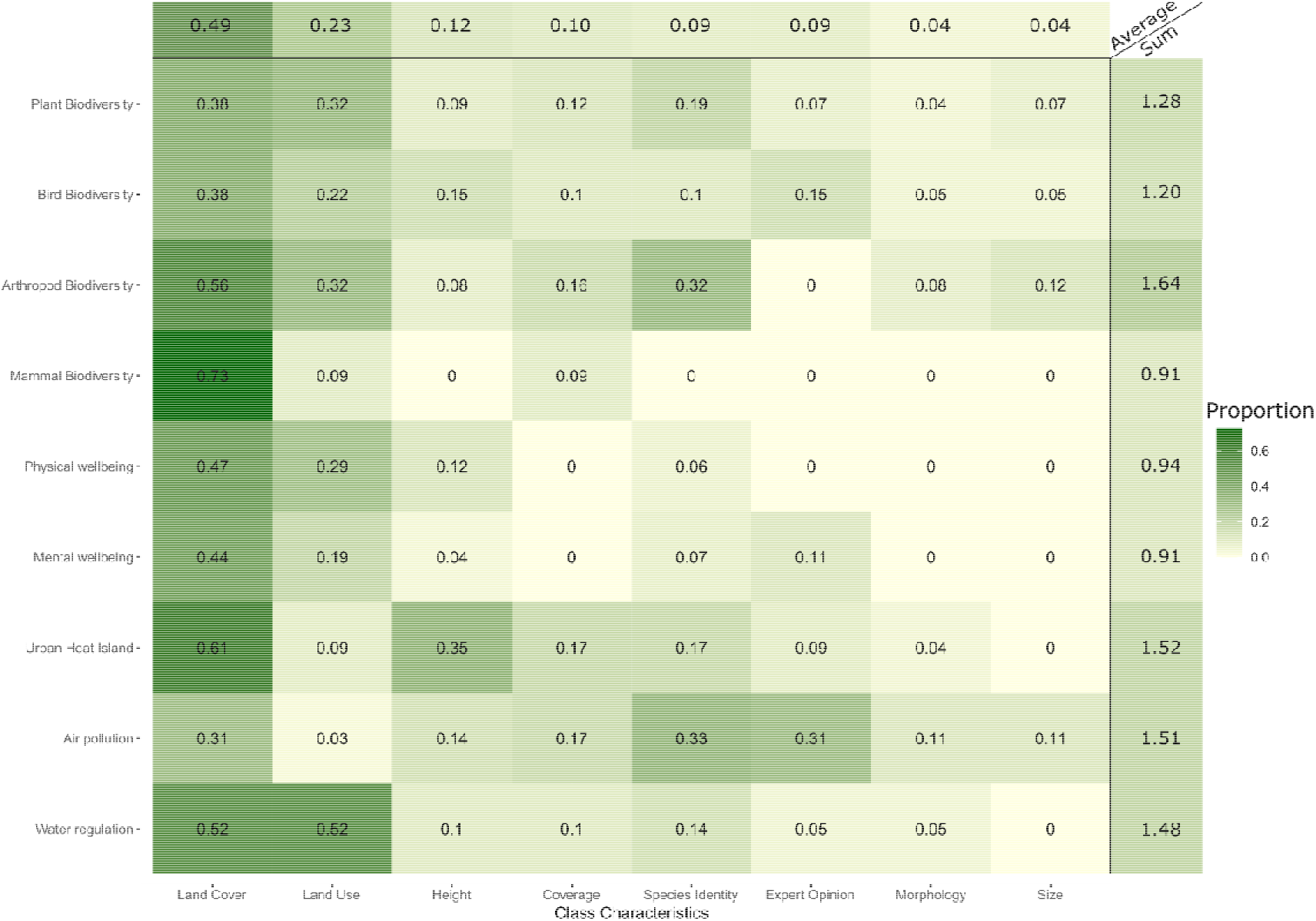
Heat map showing the proportion of characteristics used in GI classes by ES and taxon. A higher value indicates that proportionally more classes have this characteristic. Averaged and summed values are shown at end of the matrix’s columns and rows. Average values indicate use of <u>a particular</u> characteristic per ES or taxon (0-1) and summed scores indicate the average use of <u>all</u> characteristics in a ES or taxon (0-8).

### 3.4 Consolidated classification: CUGIC

We propose a classification system based on the observed themes within the class definitions and its class characteristics, obtained through literature review, consisting of a data layer with 16 LULC and a data layer with 41 vegetation classes, describing the variation of vegetation structures within the LULC classes (Figure 5, see Table 1 for definitions). We call this classification system the Consolidated Urban Green Infrastructure Classification (CUGIC).

**Figure 5.**
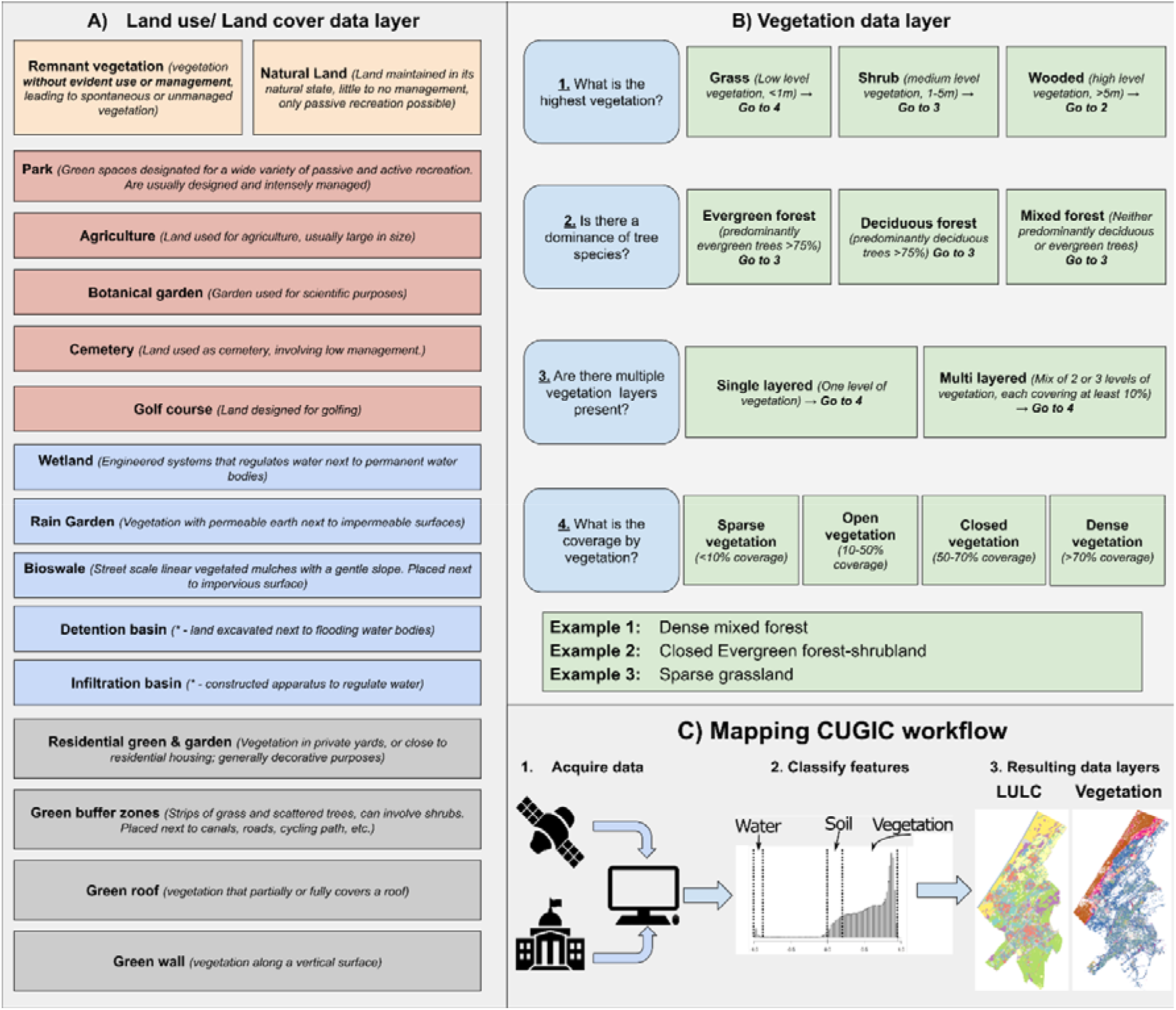
The Consolidated Urban Green Infrastructure Classification (CUGIC). Shown are all different GI classes in the consolidated classification by their data type. Data types cover: A) Land use and land cover sections which cover largely unmanaged landscapes (orange), a mix of land uses (red), water regulation (blue), and vegetation in relation to the built environment (grey). B) NDVI and height data used for vegetation structure thresholds, combining to one vegetation type (see examples 1-3). C) shows a simplified workflow of mapping the Consolidated Urban Green Infrastructure Classification (CUGIC). For further details on CUGIC, see Table 1.

**Table 1.**
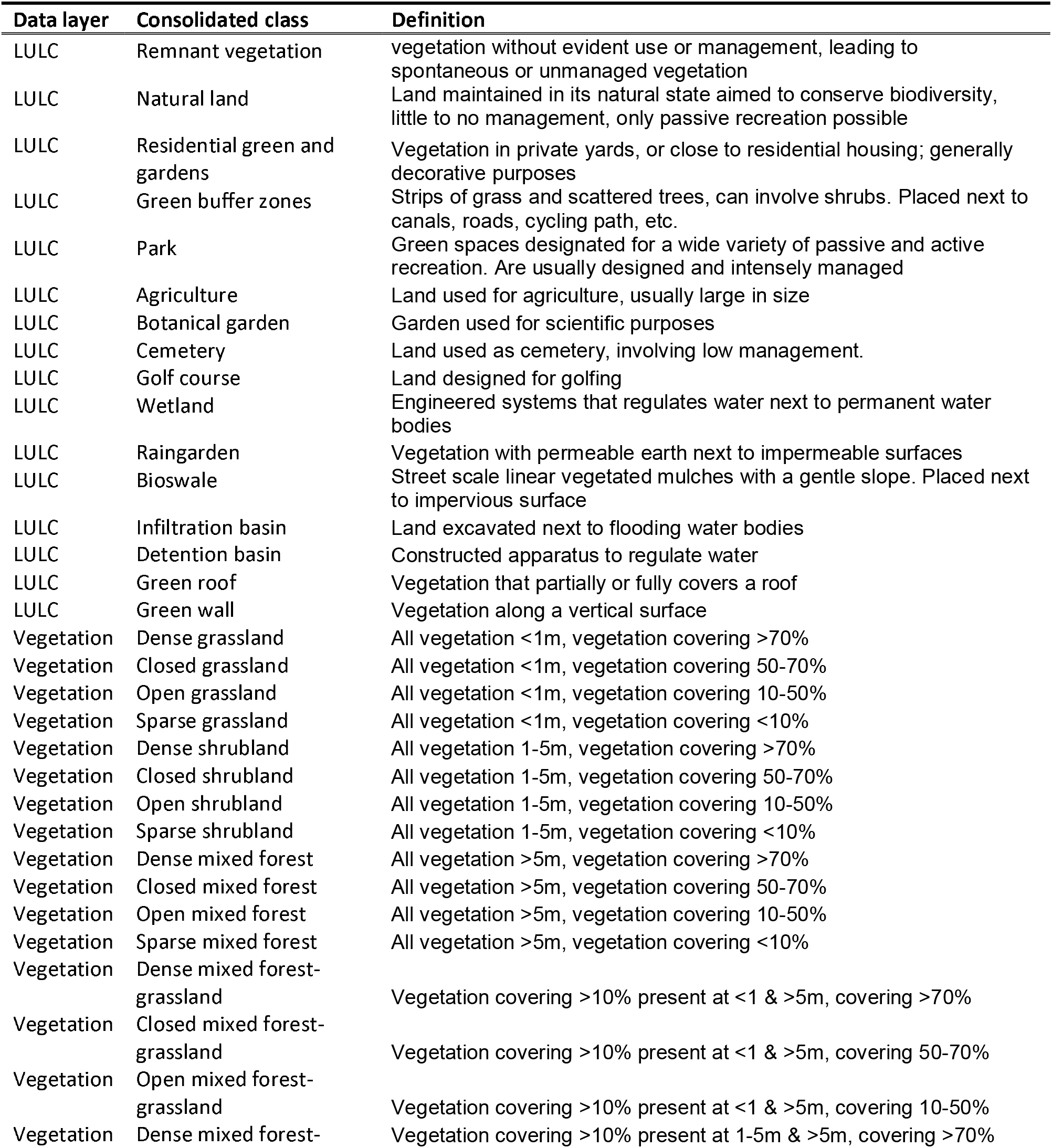
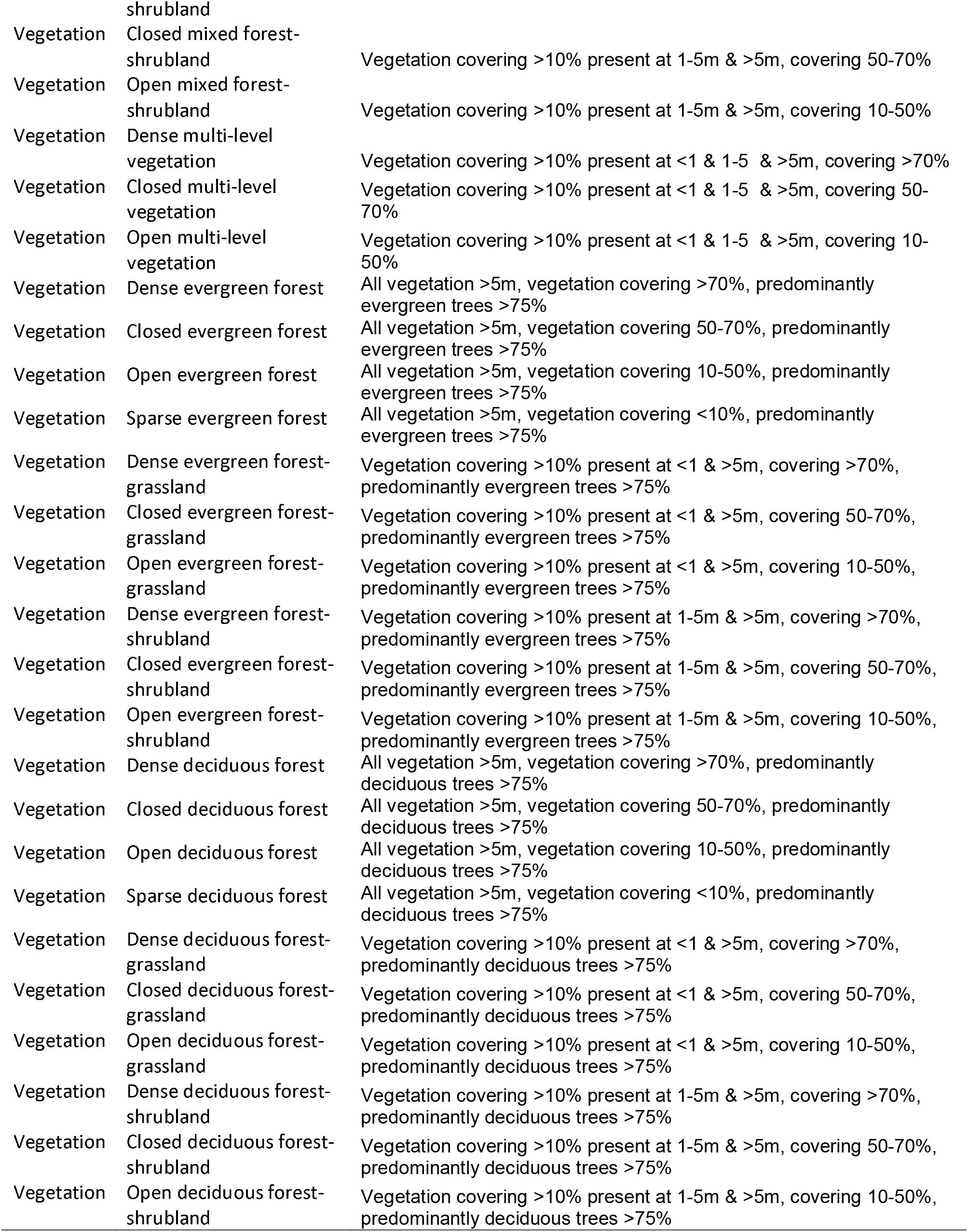
Table containing the Consolidated Urban Green Infrastructure Classification (CUGIC). Shown are all the classes from the CUGIC with their respective definitions. Note that additional lines in the table are present where the vegetational classes are solely differentiated by evergreen or deciduous trees being dominant.

CUGIC does not contain species data nor relies on expert opinions, minimizing labor to reproduce it and reducing opinion associated biases. The vegetation thresholds for height (<1, 1-5, >5) and coverage (<10%, 10-50%, 50-70%, >70%) reflect the thresholds most frequently used in GI literature. Next to height and coverage, the number of vegetation layers (as either a single or multiple layers) is included. For vegetation with trees it also takes into account dominance of evergreen or deciduous trees. See Figure 5 for a full overview of all classes and data layers included in CUGIC and its workflow.

## 4. Discussion

### 4.1 Consolidated Urban Green Infrastructure Classification - CUGIC

We presented CUGIC, the first classification scheme accommodating future multifunctional ES-biodiversity research based on literature of the past ten years (Fig. 5; Table 1). We focused on the combination between ES and biodiversity as they provide meaningful benefits to humans and nature, which are critically important in the progressively denser built urban environment (Filazzola et al., 2019; Meerow and Newell, 2019). Our novel classification is widely applicable and relevant, as (I) recent advances in remote sensing, such as open access satellite data, higher resolutions, and increased computing power, make the required data for this classification widely available (Wulder et al., 2018), (II) it should be applicable globally, allowing critical comparisons between the northern and southern hemisphere (Matsler et al., 2021), (III) the included classes are based on contemporary and well-cited ES and biodiversity literature, (IV) it answers the call for a standardized classification for interdisciplinary GI research (Bartesaghi-Koc et al., 2019; Chatzimentor et al., 2020; Matsler et al., 2021; Wang and Banzhaf, 2018), (V) allows for delineation between LULC and vegetation-driven mechanisms (Shen et al., 2021; Chaves et al., 2020), (VI) CUGIC is easy to use by mapping already often used LULC data in supplementation of indicators on vegetation. To our knowledge, this is the first Consolidated Urban GI Classification (CUGIC) of its kind based on a wide variety of literature. The presented classification is useful for both research and decision support.

### 4.2 Multifunctionality of GI classes

The class name analysis showed that across ES and biodiversity research there is a wide variety of frequently used class names (n = 49, Fig. 1). Several of these class names are unique to an ES or taxon. This suggests that those classes were designed to observe mechanisms unique to a certain aspect of GI and its related ES or taxon. For example, our analysis shows that the class rain garden is exclusively used in water regulation research, while the spontaneous landscape was only associated with plant biodiversity studies. The idea that unique GI classes could indicate unique mechanisms is well supported, as mechanisms of biodiversity functions are fundamentally different than of ES (Schwarz et al., 2017). Even within biodiversity and ES research, mechanisms that help predict ES or taxa are distinct (Beninde et al., 2015; Derkzen et al., 2015; Salmond et al., 2016). This suggests that trade-offs between ES and taxa are being reflected in unique GI classes. For example, human wellbeing is usually linked to well-managed green spaces (e.g. parks, playgrounds) (Wood et al., 2017), while native plant biodiversity metrics are generally associated to less managed green spaces (e.g. natural or spontaneous landscapes) (Threlfall et al., 2016). In this case, unique GI classes are indicators of one specific ES or taxon. For example, a bioswale, or similar water-retention structures, may mainly affect water regulation, but may not have a noticeable impact other ES or taxa. These unique mechanisms can be accounted for by incorporating mechanism-relevant classes in the typology (Bartesaghi-Koc et al., 2017). CUGIC includes a wide variety of such mechanism-relevant classes in combination with layered information on vegetation morphology and LULC, to delineate the mechanisms that drive a specific ES or taxon, allowing future research to better understand their joint relationships.

Our results also show that several GI classes are used universally by every ES or taxon, implying that they can carry out multiple functions (i.e. multifunctionality). This multifunctionality, however, has to be considered carefully as routinely used LULC-classes (Figure 4, 72%) are chosen for their superior data availability and not necessarily for their functional impacts (Andrew et al, 2015; Young et al., 2014). LULC data allows for rapid estimation of ES provisioning required for efficient urban planning (Tallis and Polasky, 2009). This rapid (but uncertain (Hazeu et al., 2011)) assessment of spatial patterns of ES may sometimes be more important than the elucidation of the relation between GI, ES, and its drivers (Dade et al., 2019). So far, only few studies explicitly mention or explain the drivers of ES (Andrew et al., 2015). To advance our mechanistic understanding of ES and biodiversity processes in urban GI, we chose to additionally include predictors of vegetation to account for intra-LULC heterogeneity in the CUGIC (Dade et al., 2019; Hazeu et al., 2011).

### 4.3 Implications

Through CUGIC, the efficiency and quality of evaluating, impacts of design, features, and long-term management can be improved to create an optimal planning of urban GI (which is currently still a topic of discussion; Sinnett et al., 2018). City planners and policy makers are increasingly interested in information on ES and biodiversity to inform their urban planning GI decisions (Grabowski et al., 2022). Additionally, multifunctionality of GI to tackle multiple urban challenges is increasingly considered (Zuniga-Teran et al., 2020). Spatial information related to GI are a core necessity for many decision support tools that quantify and map ecosystem services for urban planners (Hamel et al. 2021; van Oorschot et al., 2021). However, the diversity in GI definitions vastly increase the ambiguity in the GI concept, causing confusion among the urban planners and policy makers (Grabowski et al., 2022). A consolidated GI classification, such as CUGIC, with a clear set of GI class definitions removes the ambiguity of the terms, allowing the multiple stakeholders to communicate on an equal footing and reduce ambiguity-associated risks (Garmendia et al., 2018; Chatzimentor et al., 2020). CUGIC also allows stakeholders and planners to view urban GI from both a LULC and vegetation perspective, where the combination could lead to better informed decision making.

For the scientific realm, we designed this classification to close the gap between ES and biodiversity research in the urban environment. In particular, the relationship between ES and biodiversity requires a classification incorporating both the biophysical and social environment (Schwarz et al., 2017). While most taxa and ES have their own classification standards, the lack of alignment across the GI literature remains problematic for cross-comparisons. Therefore, CUGIC standards provide a solid foundation for future multifunctional research. This is important as synergies and trade-offs between GI types remain largely unknown, while they are required, through policy, to provide multiple benefits in an increasingly smaller available space. Usage of the CUGIC will allow for a broader understanding of the drivers of the GI-ES-biodiversity relationship. By the inclusion of a wide variety of relevant drivers, CUGIC is especially accommodating to support interdisciplinary urban GI studies.

The CUGIC also allows for better delineation of LULC and vegetation effects. We illustrate this with two cases where this classification is likely to improve our understanding of urban GI. First, human wellbeing studies usually measure green spaces through either NDVI, land cover, tree canopy or databases such as OpenStreetMap (OSM) (Labib et al., 2020). These are valid indicators of green space, yet individual use of the indicators does not allow for delineation between indicator-associated error and the effect from the vegetation or green space. For instance, the study of Ward Thompson et al. (2016) shows that the amount of green space in the neighborhood was a significant predictor of stress. While this evidence indicates that GI reduces stress, it does not elucidate what characteristic of GI reduces stress. Second, land cover classifications by experts incur bias and limit cross-comparisons. For example, forest can be defined as “area dominated by trees with height generally taller than 5m” (Tsai et al., 2018) or as “area covered predominately with trees. These areas usually contain fragments of (often degraded) forest encroached by built-up land and agricultural activities” (Wu & Kim, 2021). These definitions only overlap in relating to the factor “tree dominance”, whereas the remainder of the definitions are starkly different. In both cases, the CUGIC will allow for an improvement of the mechanistic understanding and standardization of the methodological approaches. Mainly, the CUGIC provides standardized definitions of GI aiming to reduce the human error inherent to expert opinion based metrics, allowing for cross comparison and increased interpretability of the study area (Eriksen et al., 2019).

Using CUGIC is simply mapping the LULC with indicators on vegetation structure. In addition, CUGIC can be adjusted to fit to a wide and diverse set of scenarios related to multifunctional or single function ES-biodiversity research. Although some limitations are fundamental to using classifications, such as I) containing classes irrelevant to the study-mechanism, II) the exclusion of intra-class variation, and III) being static through exclusion of novel important classes, these can be tackled with slight adjustments to CUGIC. For instance, in case that not all ES or taxa are included, the user may opt for a parsimonious solution by reducing the number of classes through discarding or combining classes (Danasingh et al., 2020). Even though this lessens the consistency among use cases, it can improve model accuracy, and using a reduction or combination of well-defined standardized classes is a significant step forward from the contemporary inconsistent usage of GI classes (Andrew et al., 2015; Bartesaghi-Koc et al., 2019). Alternatively, for some cases, it may be important to include more subtle variation in vegetation. Here, we suggest to decompose the vegetation classes by using their original numerical predictors (height, fractional vegetation cover, and number of vegetation layers). This allows for the inclusion of more accurate GI data, although it reduces cross-comparison with CUGIC. This can be off-set by extensively describing the distributions of the predictors. Moreover, considering the novelty of the GI framework, future work may develop important new GI classes. In particular, the high NIR found for birds, plants, mammals, mental and physical wellbeing indicate that a variety of novel GI is frequently tested, while few are used consistently. This may indicate that highly predictive GI classes that are used universally within a research field have not yet been created. When future research necessitates such classes, they can be easily incorporated in our proposed classification in a position where the class is mutually exclusive with other classes. Such new classes may relate to size, morphology, or species identity, which are currently absent in classifications while they are known to be drivers of biodiversity and ES (Beninde et al., 2015; Andrew et al., 2015). As a wider array of ES or taxa is considered, future research could add-on new GI classes that are relevant drivers of the respective ES or taxa to the proposed classification. Through reducing, decomposing, or including novel classes we present an easily applicable and flexible consolidated GI classification.

## 5. Conclusions

We provided a Consolidated Urban Green Infrastructure Classification (CUGIC) that is grounded in a broad urban ES and biodiversity literature. The classification can be used as tool to further unravel the complex relationship GI has with ES and biodiversity. In particular, it can be applied across the globe, it makes use of existing literature, and it explicitly addresses previous research concerns for the need of synthesized GI definitions (Shen et al, 2021; Chatzimentor et al., 2020; Grabowski et al., 2022). CUGIC is the first freely available, standardized method aimed to facilitate research that couples ES and biodiversity in the urban context. It provides unprecedented opportunities to research synergies and trade-offs within and between ES and biodiversity.

Finally, we thoroughly analyzed the GI class usage and we showed that: (i) GI classes that uniquely link to specific ES and biodiversity taxon may indicate process-driven GI classification, (ii) universal GI classes capture a variety of ES and biodiversity mechanisms through their diversity of definitions, (iii) usage of GI class characteristics is similar across ES and taxa while the mechanisms that explain ES and biodiversity are distinct. We argue that these results indicate that past research was mainly dominated by data availability and that, in line with other studies, contemporary research should aim to gain a mechanistic understanding of GI, ES and biodiversity in the urban context. Our CUGIC classification provides a novel opportunity to study the features and multiple outcomes of ES and biodiversity in urban GI, through capturing a broad variety of GI characteristics.

## 6. Data accessibility

Collected data, search queries and code used for this study have been made permanently and publicly available on the Mendeley Data repository at TBA.

## List of appendices

Appendix 1. Green Infrastructure Classifications

Appendix 2. Method for calculating saturation

Appendix 3. Search query’s used on Web of Science

Appendix 4. List of included studies

Appendix 5. Analysis of grey and blue infrastructure

Appendix 6. Details on class characteristics binary scoring

Appendix 7. Themes to CUGIC

