## Appendix 1 for "CUGIC: The Consolidated Urban Green Infrastructure Classification for assessing ecosystem services and biodiversity"

Urban Vegetation Structure Types (UVST; Lehmann et al., 2014) is quite extensive, considering a total of 57 categories. While this classification is elaborate, producing it is laborious, requiring a digital urban biotope map, which is made using aerial images and additional on-site inspection. Our Consolidated Urban Green Infrastructure Classification (CUGIC) is reproducible without on-site inspection, which significantly reduces the workload of creating it. Additionally, the UVST vegetation categories contain a significant amount of bias as they require visual inspection and use loosely defined terms, being: sparse, of low density, in groups, in rows, and dense. Here the CUGIC allows for standardisation of GI typology across studies by defining all thresholds.

The High Ecological Resolution Classification for Urban Landscapes and Environmental Systems (HERCULES; Cadenasso et al., 2007) classification is based on patches and results in 6 data layers for every patch it considers. This classification does not primarily quantify green infrastructure (Coarse vegetation, Fine vegetation), but mainly grey infrastructure (Bare soil, Pavement, Building proportion, Building type). Our CUGIC aim is to further elaborate on GI types rather than explaining ES through grey infrastructure. Here, we work largely with the same indicators, being coverage and height of vegetation, yet vastly expand on it by considering land-use which is important to biodiversity.

Green Infrastructure Typology (GIT; Bartesaghi-Koc et al., 2017; Bartsaghi-Koc et al., 2019) does not consider land-use and provides only five categories (green open space, water bodies, tree canopy, green roofs, vertical greenery). In the subsequent articles it was expanded combining the GIT with other classification criteria such as those from the UVST, Local Climate Zones (LCZ; Stewart et al., 2013), and HERCULES, accounting for functional, structural, and configuration factors, yet still lacking land-use, which seems to be paramount to many biodiversity taxa.

Moreover, while these schemes provide hard categories, we presented the CUGIC to be flexible by allowing decomposing, reducing, and implementation of new classes to fit to users’ needs while still presenting the necessary data for cross-comparability between studies.

*Literature*

Bartesaghi-Koc, C., Osmond, P., & Peters, A. (2019). Mapping and classifying green infrastructure typologies for climate-related studies based on remote sensing data. *Urban Forestry and Urban Greening*, *37*(July 2017), 154–167. https://doi.org/10.1016/j.ufug.2018.11.008

Bartesaghi Koc, C., Osmond, P., & Peters, A. (2017). Towards a comprehensive green infrastructure typology: a systematic review of approaches, methods and typologies. *Urban Ecosystems*, *20*(1), 15–35. <https://doi.org/10.1007/s11252-016-0578-5>

Cadenasso, M.L., Pickett, S.T.A., Schwarz, K., 2007. Spatial heterogeneity in urban ecosystems: reconceptualizing land cover and a framework for classification. Frontiers in Ecology and the Environment 5, 80–88.

Lehmann, I., Mathey, J., Rößler, S., Bräuer, A., Goldberg, V., 2014. Urban vegetation structure types as a methodological approach for identifying ecosystem services – Application to the analysis of micro-climatic effects. Ecological Indicators 42, 58–72. https://doi.org/10.1016/j.ecolind.2014.02.036

Stewart, I.D., Oke, T.R., Krayenhoff, E.S., 2014. Evaluation of the ‘local climate zone’ scheme using temperature observations and model simulations. International Journal of Climatology 34, 1062–1080. https://doi.org/10.1002/joc.3746
