## Appendix 2 for "CUGIC: The Consolidated Urban Green Infrastructure Classification for assessing ecosystem services and biodiversity"

***Appendix 2. Method of calculation saturation in detail***

Through measure thematic saturation we aim to report clearly the confidence in our results (Guest et al., 2020). By calculating the saturation by each taxon or ES, we get estimate the New Information Ratio (NIR) which indicates the chance of finding new information. Research papers were included if they contained at least two different classes of GI, were used to predict one ES or biodiversity taxon, and were located in the urban environment. Importantly, this excludes studies that solely use NDVI or tree canopy as a predictor, and it excludes studies incorporating multiple ES or biodiversity taxa.

To calculate saturation we used the names of the GI classes as data. The NIR then indicates the chance that incorporating an additional paper will result in a new GI class. Though many GI class names differed slightly between papers artificially increasing the NIR while the actual classes were very similar. Therefore we chose to assign a common name to each GI class and add additional GI class names if they were near-identical or meant the same. For example, classes such as “park” and “urban park” are considered as one common class name, “park”. This resulted in a large amount of common class names (n = 564) of which only 456 were used more than once, substantially deflating the NIR found for each taxon or ES.

Our chosen parameters for measure saturation was a base size of five, run length of three and a NIR of 10%. In practice, this meant that for each ES or taxon five papers were selected and the number of unique common classes were noted (base size). Next, three more papers were selected (run length) and their number of new unique common class were divided by the number of unique common classes found in the base size, resulting in the NIR. Additional papers were collected until the final three analyzed papers resulted in a NIR of equal or less than 10%. All papers related to a specific ES or taxon were investigated if the NIR stayed above the set 10% NIR threshold.
