## Appendix 3 for "CUGIC: The Consolidated Urban Green Infrastructure Classification for assessing ecosystem services and biodiversity"

***Appendix 6. Table with search queries for ES and biodiversity.*** *Both green infrastructure and urban green space were used in the first term to increase yield while incorporating near identical keywords (Matsler et al., 2021). Among the nine search queries only the second search term is different between them. This term is related to the specific ES or taxon. The TS operator searchers for “Title-Abstract-Keywords-Author Keywords”.*

| **Ecosystem service or biodiversity taxon** | **Search term** |
| --- | --- |
| Heat reduction | TS=(green infrastructure OR urban green space) AND TS=(urban heat island) AND TS=(class* OR typ*) |
| Water regulation | TS=(green infrastructure OR urban green space) AND TS=(water) AND TS=(class* OR typ*) |
| Air purification | TS=(green infrastructure OR urban green space) AND TS=(air pollution) AND TS=(class* OR typ*) |
| Mental health | TS=(green infrastructure OR urban green space) AND TS=(health AND mental) AND TS=(class* OR typ*) |
| Physical health | TS=(green infrastructure OR urban green space) AND TS=(health AND physical) AND TS=(class* OR typ*) |
| Bird biodiversity | TS=(green infrastructure OR urban green space) AND TS=(biodiversity AND bird*) AND TS=(class* OR typ*) |
| Plant biodiversity | TS=(green infrastructure OR urban green space) AND TS=(biodiversity AND plant*) AND TS=(class* OR typ*) |
| Arthropod biodiversity | TS=(green infrastructure OR urban green space) AND TS=(biodiversity AND arthropod*) AND TS=(class* OR typ*) |
| Mammal biodiversity | TS=(green infrastructure OR urban green space) AND TS=(biodiversity AND mammal*) AND TS=(class* OR typ*) |
