## Appendix 4 for "CUGIC: The Consolidated Urban Green Infrastructure Classification for assessing ecosystem services and biodiversity"

**Appendix 4. Studies included in the analysis.**

Abhijith, K. V., Kumar, P., 2021. Evaluation of respiratory deposition doses in the presence of green infrastructure. Air Quality, Atmosphere and Health 14, 911–924. https://doi.org/10.1007/s11869-021-00989-w

Abhijith, K. V., Kumar, P., 2019. Field investigations for evaluating green infrastructure effects on air quality in open-road conditions. Atmospheric Environment 201, 132–147. https://doi.org/10.1016/j.atmosenv.2018.12.036

Abhijith, K. V., Kumar, P., Gallagher, J., McNabola, A., Baldauf, R., Pilla, F., Broderick, B., Di Sabatino, S., Pulvirenti, B., 2017. Air pollution abatement performances of green infrastructure in open road and built-up street canyon environments – A review. Atmospheric Environment 162, 71–86. https://doi.org/10.1016/j.atmosenv.2017.05.014

Anderson, E.C., Minor, E.S., 2019. Assessing social and biophysical drivers of spontaneous plant diversity and structure in urban vacant lots. Science of the Total Environment 653, 1272–1281. https://doi.org/10.1016/j.scitotenv.2018.11.006

Anderson, V., Gough, W.A., 2020. Evaluating the potential of nature-based solutions to reduce ozone, nitrogen dioxide, and carbon dioxide through a multi-type green infrastructure study in Ontario, Canada. City and Environment Interactions 6, 100043. https://doi.org/10.1016/j.cacint.2020.100043

Appalasamy, M., Ramdhani, S., Sershen, 2020. Aliens in the city: Towards identifying non-indigenous floristic hotspots within an urban matrix. Flora: Morphology, Distribution, Functional Ecology of Plants 269, 151631. https://doi.org/10.1016/j.flora.2020.151631

Ariza, S.L.J., Martínez, J.A., Muñoz, A.F., Quijano, J.P., Rodríguez, J.P., Camacho, L.A., Díaz-Granados, M., 2019. A multicriteria planning framework to locate and select sustainable urban drainage systems (SUDS) in consolidated urban areas. Sustainability (Switzerland) 11, 2312. https://doi.org/10.3390/su11082312

Astell-Burt, T., Feng, X., 2019. Association of Urban Green Space with Mental Health and General Health among Adults in Australia. JAMA Network Open 2, e198209. https://doi.org/10.1001/jamanetworkopen.2019.8209

Astell-Burt, T., Navakatikyan, M.A., Feng, X., 2020. Urban green space, tree canopy and 11-year risk of dementia in a cohort of 109,688 Australians. Environment International 145, 106102. https://doi.org/10.1016/j.envint.2020.106102

Astell-Burt, T., Navakatikyan, M.A., Walsan, R., Davis, W., Figtree, G., Arnolda, L., Feng, X., 2021. Green space and cardiovascular health in people with type 2 diabetes. Health and Place 69, 102554. https://doi.org/10.1016/j.healthplace.2021.102554

Balbi, M., Petit, E.J., Croci, S., Nabucet, J., Georges, R., Madec, L., Ernoult, A., 2019. Title: Ecological relevance of least cost path analysis: An easy implementation method for landscape urban planning. Journal of Environmental Management 244, 61–68. https://doi.org/10.1016/j.jenvman.2019.04.124

Barnes, M.R., Donahue, M.L., Keeler, B.L., Shorb, C.M., Mohtadi, T.Z., Shelby, L.J., 2019. Characterizing nature and participant experience in studies of nature exposure for positive mental health: An integrative review. Frontiers in Psychology 9. https://doi.org/10.3389/fpsyg.2018.02617

Bartesaghi Koc, C., Osmond, P., Peters, A., 2018. Evaluating the cooling effects of green infrastructure: A systematic review of methods, indicators and data sources. Solar Energy 166, 486–508. https://doi.org/10.1016/j.solener.2018.03.008

Barwise, Y., Kumar, P., 2020. Designing vegetation barriers for urban air pollution abatement: a practical review for appropriate plant species selection. npj Climate and Atmospheric Science 3, 12. https://doi.org/10.1038/s41612-020-0115-3

Battisti, L., Pille, L., Wachtel, T., Larcher, F., Säumel, I., 2019. Residential greenery: State of the art and health-related ecosystem services and disservices in the city of Berlin. Sustainability (Switzerland) 11, 1815. https://doi.org/10.3390/su11061815

Belcher, R.N., Sadanandan, K.R., Goh, E.R., Chan, J.Y., Menz, S., Schroepfer, T., 2019. Vegetation on and around large-scale buildings positively influences native tropical bird abundance and bird species richness. Urban Ecosystems 22, 213–225. https://doi.org/10.1007/s11252-018-0808-0

Bennett, A.B., Gratton, C., 2012. Local and landscape scale variables impact parasitoid assemblages across an urbanization gradient. Landscape and Urban Planning 104, 26–33. https://doi.org/10.1016/j.landurbplan.2011.09.007

Bokaie, M., Zarkesh, M.K., Arasteh, P.D., Hosseini, A., 2016. Assessment of Urban Heat Island based on the relationship between land surface temperature and Land Use/ Land Cover in Tehran. Sustainable Cities and Society 23, 94–104. https://doi.org/10.1016/j.scs.2016.03.009

Bonthoux, S., Voisin, L., Bouché-Pillon, S., Chollet, S., 2019. More than weeds: Spontaneous vegetation in streets as a neglected element of urban biodiversity. Landscape and Urban Planning 185, 163–172. https://doi.org/10.1016/j.landurbplan.2019.02.009

Borysiak, J., Mizgajski, A., Speak, A., 2017. Floral biodiversity of allotment gardens and its contribution to urban green infrastructure. Urban Ecosystems 20, 323–335. https://doi.org/10.1007/s11252-016-0595-4

Bottalico, F., Travaglini, D., Chirici, G., Garfì, V., Giannetti, F., De Marco, A., Fares, S., Marchetti, M., Nocentini, S., Paoletti, E., Salbitano, F., Sanesi, G., 2017. A spatially-explicit method to assess the dry deposition of air pollution by urban forests in the city of Florence, Italy. Urban Forestry and Urban Greening 27, 221–234. https://doi.org/10.1016/j.ufug.2017.08.013

Cai, L., Zhuang, M., Ren, Y., 2020. A landscape scale study in Southeast China investigating the effects of varied green space types on atmospheric PM2.5 in mid-winter. Urban Forestry and Urban Greening 49, 126607. https://doi.org/10.1016/j.ufug.2020.126607

Carbó-Ramírez, P., Zuria, I., 2011. The value of small urban greenspaces for birds in a Mexican city. Landscape and Urban Planning 100, 213–222. https://doi.org/10.1016/j.landurbplan.2010.12.008

Casal-Campos, A., Sadr, S.M.K., Fu, G., Butler, D., 2018. Reliable, Resilient and Sustainable Urban Drainage Systems: An Analysis of Robustness under Deep Uncertainty. Environmental Science and Technology 52, 9008–9021. https://doi.org/10.1021/acs.est.8b01193

Chang, C.R., Chien, H.F., Shiu, H.J., Ko, C.J., Lee, P.F., 2017. Multiscale heterogeneity within and beyond Taipei city greenspaces and their relationship with avian biodiversity. Landscape and Urban Planning 157, 138–150. https://doi.org/10.1016/j.landurbplan.2016.05.028

Chang, J., Ren, Y., Shi, Y., Zhu, Y., Ge, Y., Hong, S., Jiao, L., Lin, F., Peng, C., Mochizuki, T., Tani, A., Mu, Y., Fu, C., 2012. An inventory of biogenic volatile organic compounds for a subtropical urban-rural complex. Atmospheric Environment 56, 115–123. https://doi.org/10.1016/j.atmosenv.2012.03.053

Chen, A., Yao, X.A., Sun, R., Chen, L., 2014. Effect of urban green patterns on surface urban cool islands and its seasonal variations. Urban Forestry and Urban Greening 13, 646–654. https://doi.org/10.1016/j.ufug.2014.07.006

Chen, M., Dai, F., Yang, B., Zhu, S., 2019. Effects of urban green space morphological pattern on variation of PM2.5 concentration in the neighborhoods of five Chinese megacities. Building and Environment 158, 1–15. https://doi.org/10.1016/j.buildenv.2019.04.058

Çoban, S., Yener, Ş.D., Bayraktar, S., 2021. Woody plant composition and diversity of urban green spaces in Istanbul, Turkey. Plant Biosystems 155, 83–91. https://doi.org/10.1080/11263504.2020.1727980

Cohen, P., Potchter, O., Matzarakis, A., 2012. Daily and seasonal climatic conditions of green urban open spaces in the Mediterranean climate and their impact on human comfort. Building and Environment 51, 285–295. https://doi.org/10.1016/j.buildenv.2011.11.020

Cornell, K.L., Kight, C.R., Burdge, R.B., Gunderson, A.R., Hubbard, J.K., Jackson, A.K., LeClerc, J.E., Pitts, M.L., Swaddle, J.P., Cristol, D.A., 2011. Reproductive success of Eastern Bluebirds (Siala sialis) on suburban golf courses. Auk 128, 577–586. https://doi.org/10.1525/auk.2011.10182

Davis, A.Y., Belaire, J.A., Farfan, M.A., Milz, D., Sweeney, E.R., Loss, S.R., Minor, E.S., 2012. Green infrastructure and bird diversity across an urban socioeconomic gradient. Ecosphere 3, art105. https://doi.org/10.1890/es12-00126.1

Deng, S., Ma, J., Zhang, L., Jia, Z., Ma, L., 2019. Microclimate simulation and model optimization of the effect of roadway green space on atmospheric particulate matter. Environmental Pollution 246, 932–944. https://doi.org/10.1016/j.envpol.2018.12.026

Dennis, M., Cook, P.A., James, P., Wheater, C.P., Lindley, S.J., 2020. Relationships between health outcomes in older populations and urban green infrastructure size, quality and proximity. BMC Public Health 20, 626. https://doi.org/10.1186/s12889-020-08762-x

Dylewski, Ł., Maćkowiak, Ł., Banaszak-Cibicka, W., 2019. Are all urban green spaces a favourable habitat for pollinator communities? Bees, butterflies and hoverflies in different urban green areas. Ecological Entomology 44, 678–689. https://doi.org/10.1111/een.12744

Dzhambov, A.M., Markevych, I., Lercher, P., 2018. Greenspace seems protective of both high and low blood pressure among residents of an Alpine valley. Environment International 121, 443–452. https://doi.org/10.1016/j.envint.2018.09.044

Fischer, L.K., Rodorff, V., von der Lippe, M., Kowarik, I., 2016. Drivers of biodiversity patterns in parks of a growing South American megacity. Urban Ecosystems 19, 1231–1249. https://doi.org/10.1007/s11252-016-0537-1

Fröhlich, A., Ciach, M., 2020. Dead tree branches in urban forests and private gardens are key habitat components for woodpeckers in a city matrix. Landscape and Urban Planning 202, 103869. https://doi.org/10.1016/j.landurbplan.2020.103869

Gago, E.J., Roldan, J., Pacheco-Torres, R., Ordóñez, J., 2013. The city and urban heat islands: A review of strategies to mitigate adverse effects. Renewable and Sustainable Energy Reviews 25, 749–758. https://doi.org/10.1016/j.rser.2013.05.057

Gallo, T., Fidino, M., Lehrer, E.W., Magle, S.B., 2017. Mammal diversity and metacommunity dynamics in urban green spaces: Implications for urban wildlife conservation. Ecological Applications 27, 2330–2341. https://doi.org/10.1002/eap.1611

Gao, T., Liu, F., Wang, Y., Mu, S., Qiu, L., 2020. Reduction of atmospheric suspended particulate matter concentration and influencing factors of green space in Urban forest park. Forests 11, 950. https://doi.org/10.3390/f11090950

Gavrić, S., Leonhardt, G., Marsalek, J., Viklander, M., 2019. Processes improving urban stormwater quality in grass swales and filter strips: A review of research findings. Science of the Total Environment 669, 431–447. https://doi.org/10.1016/j.scitotenv.2019.03.072

Gittleman, M., Farmer, C.J.Q., Kremer, P., McPhearson, T., 2017. Estimating stormwater runoff for community gardens in New York City. Urban Ecosystems 20, 129–139. https://doi.org/10.1007/s11252-016-0575-8

Gonçalves, S.F., Lourenço, A.C. de P., Bueno Filho, J.S. de S., de Toledo, M.C.B., 2021. Characteristics of residential backyards that contribute to conservation and diversity of urban birds: A case study in a Southeastern Brazilian city. Urban Forestry and Urban Greening 61, 127095. https://doi.org/10.1016/j.ufug.2021.127095

Gong, C., Chen, J., Yu, S., 2013. Biotic homogenization and differentiation of the flora in artificial and near-natural habitats across urban green spaces. Landscape and Urban Planning 120, 158–169. https://doi.org/10.1016/j.landurbplan.2013.08.006

Grafius, D.R., Corstanje, R., Siriwardena, G.M., Plummer, K.E., Harris, J.A., 2017. A bird’s eye view: using circuit theory to study urban landscape connectivity for birds. Landscape Ecology 32, 1771–1787. https://doi.org/10.1007/s10980-017-0548-1

Gunnell, K., Mulligan, M., Francis, R.A., Hole, D.G., 2019. Evaluating natural infrastructure for flood management within the watersheds of selected global cities. Science of the Total Environment 670, 411–424. https://doi.org/10.1016/j.scitotenv.2019.03.212

Helen, Jarzebski, M.P., Gasparatos, A., 2019. Land use change, carbon stocks and tree species diversity in green spaces of a secondary city in Myanmar, Pyin Oo Lwin. PLoS ONE 14, e0225331. https://doi.org/10.1371/journal.pone.0225331

Houlden, V., Porto de Albuquerque, J., Weich, S., Jarvis, S., 2021. Does nature make us happier? A spatial error model of greenspace types and mental wellbeing. Environment and Planning B: Urban Analytics and City Science 48, 655–670. https://doi.org/10.1177/2399808319887395

Huang, F., Zhang, Y., Lou, Y., Li, X., Zhang, T., Yu, H., Yuan, C., Tong, Q., Qi, F., Shao, F., 2021. Characterization, Sources and Excessive Cancer Risk of PM2.5-Bound Polycyclic Aromatic Hydrocarbons in Different Green Spaces in Lin’an, Hangzhou, China. Bulletin of Environmental Contamination and Toxicology 107, 519–529. https://doi.org/10.1007/s00128-021-03304-6

Huang, Q., Yang, M., Jane, H. ann, Li, S., Bauer, N., 2020. Trees, grass, or concrete? The effects of different types of environments on stress reduction. Landscape and Urban Planning 193, 103654. https://doi.org/10.1016/j.landurbplan.2019.103654

Hüse, B., Szabó, S., Deák, B., Tóthmérész, B., 2016. Mapping an ecological network of green habitat patches and their role in maintaining urban biodiversity in and around Debrecen city (Eastern Hungary). Land Use Policy 57, 574–581. https://doi.org/10.1016/j.landusepol.2016.06.026

Ibáñez-Álamo, J.D., Morelli, F., Benedetti, Y., Rubio, E., Jokimäki, J., Pérez-Contreras, T., Sprau, P., Suhonen, J., Tryjanowski, P., Kaisanlahti-Jokimäki, M.L., Møller, A.P., Díaz, M., 2020. Biodiversity within the city: Effects of land sharing and land sparing urban development on avian diversity. Science of the Total Environment 707, 135477. https://doi.org/10.1016/j.scitotenv.2019.135477

Itani, M., Al Zein, M., Nasralla, N., Talhouk, S.N., 2020. Biodiversity conservation in cities: Defining habitat analogues for plant species of conservation interest. PLoS ONE 15, e0220355. https://doi.org/10.1371/journal.pone.0220355

Jaganmohan, M., Knapp, S., Buchmann, C.M., Schwarz, N., 2016. The Bigger, the Better? The Influence of Urban Green Space Design on Cooling Effects for Residential Areas. Journal of Environmental Quality 45, 134–145. https://doi.org/10.2134/jeq2015.01.0062

Kazemi, F., Beecham, S., Gibbs, J., 2011. Streetscape biodiversity and the role of bioretention swales in an Australian urban environment. Landscape and Urban Planning 101, 139–148. https://doi.org/10.1016/j.landurbplan.2011.02.006

Keten, A., Eroglu, E., Kaya, S., Anderson, J.T., 2020. Bird diversity along a riparian corridor in a moderate urban landscape. Ecological Indicators 118, 106751. https://doi.org/10.1016/j.ecolind.2020.106751

Korányi, D., Gallé, R., Donkó, B., Chamberlain, D.E., Batáry, P., 2021. Urbanization does not affect green space bird species richness in a mid-sized city. Urban Ecosystems 24, 789–800. https://doi.org/10.1007/s11252-020-01083-2

Kuang, W., Liu, Y., Dou, Y., Chi, W., Chen, G., Gao, C., Yang, T., Liu, J., Zhang, R., 2015. What are hot and what are not in an urban landscape: quantifying and explaining the land surface temperature pattern in Beijing, China. Landscape Ecology 30, 357–373. https://doi.org/10.1007/s10980-014-0128-6

Lahoti, S., Lahoti, A., Joshi, R.K., Saito, O., 2020. Vegetation structure, species composition, and carbon sink potential of urban green spaces in Nagpur City, India. Land 9, 107. https://doi.org/10.3390/land9040107

Lanki, T., Siponen, T., Ojala, A., Korpela, K., Pennanen, A., Tiittanen, P., Tsunetsugu, Y., Kagawa, T., Tyrväinen, L., 2017. Acute effects of visits to urban green environments on cardiovascular physiology in women: A field experiment. Environmental Research 159, 176–185. https://doi.org/10.1016/j.envres.2017.07.039

Leng, H., Li, S., Yan, S., An, X., 2020. Exploring the relationship between green space in a neighbourhood and cardiovascular health in the winter city of China: A study using a health survey for harbin. International Journal of Environmental Research and Public Health 17, 513. https://doi.org/10.3390/ijerph17020513

Li, X.P., Fan, S.X., Hao, P.Y., Dong, L., 2019. Temporal variations of spontaneous plants colonizing in different type of planted vegetation-a case of Beijing Olympic Forest Park. Urban Forestry and Urban Greening 46, 126459. https://doi.org/10.1016/j.ufug.2019.126459

Lintott, P.R., Barlow, K., Bunnefeld, N., Briggs, P., Gajas Roig, C., Park, K.J., 2016. Differential responses of cryptic bat species to the urban landscape. Ecology and Evolution 6, 2044–2052. https://doi.org/10.1002/ece3.1996

Lintott, P.R., Bunnefeld, N., Minderman, J., Fuentes-Montemayor, E., Mayhew, R.J., Olley, L., Park, K.J., 2015. Differential responses to woodland character and landscape context by cryptic bats in urban environments. PLoS ONE 10, e0126850. https://doi.org/10.1371/journal.pone.0126850

Lobaccaro, G., Acero, J.A., 2015. Comparative analysis of green actions to improve outdoor thermal comfort inside typical urban street canyons. Urban Climate 14, 251–267. https://doi.org/10.1016/j.uclim.2015.10.002

Luan, B., Yin, R., Xu, P., Wang, X., Yang, X., Zhang, L., Tang, X., 2019. Evaluating Green Stormwater Infrastructure strategies efficiencies in a rapidly urbanizing catchment using SWMM-based TOPSIS. Journal of Cleaner Production 223, 680–691. https://doi.org/10.1016/j.jclepro.2019.03.028

Lucas, W.C., Sample, D.J., 2015. Reducing combined sewer overflows by using outlet controls for Green Stormwater Infrastructure: Case study in Richmond, Virginia. Journal of Hydrology 520, 473–488. https://doi.org/10.1016/j.jhydrol.2014.10.029

Maes, M.J.A., Pirani, M., Booth, E.R., Shen, C., Milligan, B., Jones, K.E., Toledano, M.B., 2021. Benefit of woodland and other natural environments for adolescents’ cognition and mental health. Nature Sustainability 4, 851–858. https://doi.org/10.1038/s41893-021-00751-1

Manes, F., Marando, F., Capotorti, G., Blasi, C., Salvatori, E., Fusaro, L., Ciancarella, L., Mircea, M., Marchetti, M., Chirici, G., Munafò, M., 2016. Regulating Ecosystem Services of forests in ten Italian Metropolitan Cities: Air quality improvement by PM10 and O3 removal. Ecological Indicators 67, 425–440. https://doi.org/10.1016/j.ecolind.2016.03.009

Maragno, D., Gaglio, M., Robbi, M., Appiotti, F., Fano, E.A., Gissi, E., 2018. Fine-scale analysis of urban flooding reduction from green infrastructure: An ecosystem services approach for the management of water flows. Ecological Modelling 386, 1–10. https://doi.org/10.1016/j.ecolmodel.2018.08.002

Marselle, M.R., Irvine, K.N., Lorenzo-Arribas, A., Warber, S.L., 2015. Moving beyond green: Exploring the relationship of environment type and indicators of perceived environmental quality on emotional well-being following group walks. International Journal of Environmental Research and Public Health 12, 106–130. https://doi.org/10.3390/ijerph120100106

Marselle, M.R., Irvine, K.N., Warber, S.L., 2013. Walking for well-being: Are group walks in certain types of natural environments better for well-being than group walks in urban environments? International Journal of Environmental Research and Public Health 10, 5603–5628. https://doi.org/10.3390/ijerph10115603

Mata, L., Threlfall, C.G., Williams, N.S.G., Hahs, A.K., Malipatil, M., Stork, N.E., Livesley, S.J., 2017. Conserving herbivorous and predatory insects in urban green spaces. Scientific Reports 7, 40970. https://doi.org/10.1038/srep40970

McFarland, A.R., Larsen, L., Yeshitela, K., Engida, A.N., Love, N.G., 2019. Guide for using green infrastructure in urban environments for stormwater management. Environmental Science: Water Research and Technology 5, 643–659. https://doi.org/10.1039/c8ew00498f

McLean, P., Wilson, J.R.U., Gaertner, M., Kritzinger-Klopper, S., Richardson, D.M., 2018. The distribution and status of alien plants in a small South African town. South African Journal of Botany 117, 71–78. https://doi.org/10.1016/j.sajb.2018.02.392

Miralles-Guasch, C., Dopico, J., Delclòs-Alió, X., Knobel, P., Marquet, O., Maneja-Zaragoza, R., Schipperijn, J., Vich, G., 2019. Natural landscape, infrastructure, and health: The physical activity implications of urban green space composition among the elderly. International Journal of Environmental Research and Public Health 16, 3986. https://doi.org/10.3390/ijerph16203986

Morakinyo, T.E., Kalani, K.W.D., Dahanayake, C., Ng, E., Chow, C.L., 2017. Temperature and cooling demand reduction by green-roof types in different climates and urban densities: A co-simulation parametric study. Energy and Buildings 145, 226–237. https://doi.org/10.1016/j.enbuild.2017.03.066

Moreira, T.C.L., Polize, J.L., Brito, M., da Silva Filho, D.F., Chiavegato Filho, A.D.P., Viana, M.C., Andrade, L.H., Mauad, T., 2021. Assessing the impact of urban environment and green infrastructure on mental health: results from the São Paulo Megacity Mental Health Survey. Journal of Exposure Science and Environmental Epidemiology. https://doi.org/10.1038/s41370-021-00349-x

Moreira, T.C.L., Polizel, J.L., Santos, I. de S., Silva Filho, D.F., Bensenor, I., Lotufo, P.A., Mauad, T., 2020. Green spaces, land cover, street trees and hypertension in the megacity of são paulo. International Journal of Environmental Research and Public Health 17, 725. https://doi.org/10.3390/ijerph17030725

Morelli, F., Mikula, P., Benedetti, Y., Bussière, R., Jerzak, L., Tryjanowski, P., 2018a. Escape behaviour of birds in urban parks and cemeteries across Europe: Evidence of behavioural adaptation to human activity. Science of the Total Environment 631–632, 803–810. https://doi.org/10.1016/j.scitotenv.2018.03.118

Morelli, F., Mikula, P., Benedetti, Y., Bussière, R., Tryjanowski, P., 2018b. Cemeteries support avian diversity likewise urban parks in European cities: Assessing taxonomic, evolutionary and functional diversity. Urban Forestry and Urban Greening 36, 90–99. https://doi.org/10.1016/j.ufug.2018.10.011

Moussa, S., Kuyah, S., Kyereh, B., Tougiani, A., Mahamane, S., 2020. Diversity and structure of urban forests of Sahel cities in Niger. Urban Ecosystems 23, 851–864. https://doi.org/10.1007/s11252-020-00984-6

Müller, G., Harhoff, R., Rahe, C., Berger, K., 2018. Inner-city green space and its association with body mass index and prevalent type 2 diabetes: A cross-sectional study in an urban German city. BMJ Open 8, e019062. https://doi.org/10.1136/bmjopen-2017-019062

Nero, B.F., 2019. Woody species and trait diversity-functional relations of green spaces in Kumasi, Ghana. Urban Ecosystems 22, 593–607. https://doi.org/10.1007/s11252-019-00835-z

Nero, Bertrand Festus, Callo-Concha, D., Denich, M., 2018. Structure, diversity, and carbon stocks of the tree community of Kumasi, Ghana. Forests 9, 519. https://doi.org/10.3390/f9090519

Nero, Bertrand F., Kwapong, N.A., Jatta, R., Fatunbi, O., 2018. Tree species diversity and socioeconomic perspectives of the Urban (Food) Forest of Accra, Ghana. Sustainability (Switzerland) 10, 3417. https://doi.org/10.3390/su10103417

Ng, E., Chen, L., Wang, Y., Yuan, C., 2012. A study on the cooling effects of greening in a high-density city: An experience from Hong Kong. Building and Environment 47, 256–271. https://doi.org/10.1016/j.buildenv.2011.07.014

Nishigaki, M., Hanazato, M., Koga, C., Kondo, K., 2020. What types of greenspaces are associated with depression in urban and rural older adults?: A multilevel cross-sectional study from JAGES. International Journal of Environmental Research and Public Health 17, 1–16. https://doi.org/10.3390/ijerph17249276

Nitoslawski, S.A., Duinker, P.N., 2016. Managing tree diversity: A comparison of suburban development in two Canadian cities. Forests 7, 119. https://doi.org/10.3390/f7060119

Normandin, É., Vereecken, N.J., Buddle, C.M., Fournier, V., 2017. Taxonomic and functional trait diversity of wild bees in different urban settings. PeerJ 2017, e3051. https://doi.org/10.7717/peerj.3051

Norton, B.A., Coutts, A.M., Livesley, S.J., Harris, R.J., Hunter, A.M., Williams, N.S.G., 2015. Planning for cooler cities: A framework to prioritise green infrastructure to mitigate high temperatures in urban landscapes. Landscape and Urban Planning 134, 127–138. https://doi.org/10.1016/j.landurbplan.2014.10.018

Paciência, I., Cavaleiro Rufo, J., Ribeiro, A.I., Mendes, F.C., Farraia, M., Silva, D., Delgado, L., Moreira, A., 2021. Association between the density and type of trees around urban schools and exhaled nitric oxide levels in schoolchildren. European Annals of Allergy and Clinical Immunology 53, 29–36. https://doi.org/10.23822/EurAnnACI.1764-1489.162

Parsons, A.W., Forrester, T., Baker-Whatton, M.C., McShea, W.J., Rota, C.T., Schuttler, S.G., Millspaugh, J.J., Kays, R., 2018. Mammal communities are larger and more diverse in moderately developed areas. eLife 7. https://doi.org/10.7554/eLife.38012

Pellissier, V., Cohen, M., Boulay, A., Clergeau, P., 2012. Birds are also sensitive to landscape composition and configuration within the city centre. Landscape and Urban Planning 104, 181–188. https://doi.org/10.1016/j.landurbplan.2011.10.011

Peng, J., Xie, P., Liu, Y., Ma, J., 2016. Urban thermal environment dynamics and associated landscape pattern factors: A case study in the Beijing metropolitan region. Remote Sensing of Environment 173, 145–155. https://doi.org/10.1016/j.rse.2015.11.027

Perales-Momparler, S., Andrés-Doménech, I., Hernández-Crespo, C., Vallés-Morán, F., Martín, M., Escuder-Bueno, I., Andreu, J., 2017. The role of monitoring sustainable drainage systems for promoting transition towards regenerative urban built environments: a case study in the Valencian region, Spain. Journal of Cleaner Production 163, S113–S124. https://doi.org/10.1016/j.jclepro.2016.05.153

Philpott, S.M., Cotton, J., Bichier, P., Friedrich, R.L., Moorhead, L.C., Uno, S., Valdez, M., 2014. Local and landscape drivers of arthropod abundance, richness, and trophic composition in urban habitats. Urban Ecosystems 17, 513–532. https://doi.org/10.1007/s11252-013-0333-0

Planchuelo, G., Kowarik, I., von der Lippe, M., 2020. Plant traits, biotopes and urbanization dynamics explain the survival of endangered urban plant populations. Journal of Applied Ecology 57, 1581–1592. https://doi.org/10.1111/1365-2664.13661

Qiu, L., Liu, F., Zhang, X., Gao, T., 2018. The reducing effect of green spaces with different vegetation structure on atmospheric particulate matter concentration in BaoJi City, China. Atmosphere 9, 332. https://doi.org/10.3390/atmos9090332

Qiu, L., Zhu, L., Chang, P., Wang, J., Fan, J., Gao, T., 2021. Is urban spontaneous vegetation rich in species and has potential for exploitation? - A case study in Baoji, China. Plant Biosystems 155, 42–53. https://doi.org/10.1080/11263504.2019.1701125

Rasul, A., Balzter, H., Smith, C., 2015. Spatial variation of the daytime Surface Urban Cool Island during the dry season in Erbil, Iraqi Kurdistan, from Landsat 8. Urban Climate 14, 176–186. https://doi.org/10.1016/j.uclim.2015.09.001

Rico-Silva, J.F., Cruz-Trujillo, E.J., Colorado Z, G.J., 2021. Influence of environmental factors on bird diversity in greenspaces in an Amazonian city. Urban Ecosystems 24, 365–374. https://doi.org/10.1007/s11252-020-01042-x

Riley, C.B., Herms, D.A., Gardiner, M.M., 2018. Exotic trees contribute to urban forest diversity and ecosystem services in inner-city Cleveland, OH. Urban Forestry and Urban Greening 29, 367–376. https://doi.org/10.1016/j.ufug.2017.01.004

Roberts, H., van Lissa, C., Hagedoorn, P., Kellar, I., Helbich, M., 2019. The effect of short-term exposure to the natural environment on depressive mood: A systematic review and meta-analysis. Environmental Research 177, 108606. https://doi.org/10.1016/j.envres.2019.108606

Rudolph, M., Velbert, F., Schwenzfeier, S., Kleinebecker, T., Klaus, V.H., 2017. Patterns and potentials of plant species richness in high- and low-maintenance urban grasslands. Applied Vegetation Science 20, 18–27. https://doi.org/10.1111/avsc.12267

Rupprecht, C.D.D., Byrne, J.A., 2014. Informal urban green-space: Comparison of quantity and characteristics in Brisbane, Australia and Sapporo, Japan. PLoS ONE 9, e99784. https://doi.org/10.1371/journal.pone.0099784

Schwartz, A.J., Dodds, P.S., O’Neil-Dunne, J.P.M., Danforth, C.M., Ricketts, T.H., 2019. Visitors to urban greenspace have higher sentiment and lower negativity on Twitter. People and Nature 1, 476–485. https://doi.org/10.1002/pan3.10045

Sehrt, M., Bossdorf, O., Freitag, M., Bucharova, A., 2020. Less is more! Rapid increase in plant species richness after reduced mowing in urban grasslands. Basic and Applied Ecology 42, 47–53. https://doi.org/10.1016/j.baae.2019.10.008

Shih, W.Y., 2018. Bird diversity of greenspaces in the densely developed city centre of Taipei. Urban Ecosystems 21, 379–393. https://doi.org/10.1007/s11252-017-0720-z

Silva, C.P., García, C.E., Estay, S.A., Barbosa, O., Chapman, M.G., 2015. Bird richness and abundance in response to urban form in a Latin American City: Valdivia, Chile as a Case Study. PLoS ONE 10, e0138120. https://doi.org/10.1371/journal.pone.0138120

Smith, A.C., Francis, C.M., Fahrig, L., 2014. Similar effects of residential and non-residential vegetation on bird diversity in suburban neighbourhoods. Urban Ecosystems 17, 27–44. https://doi.org/10.1007/s11252-013-0301-8

Steenberg, J.W.N., 2018. People or place? An exploration of social and ecological drivers of urban forest species composition. Urban Ecosystems 21, 887–901. https://doi.org/10.1007/s11252-018-0764-8

Sun, R., Chen, L., 2017. Effects of green space dynamics on urban heat islands: Mitigation and diversification. Ecosystem Services 23, 38–46. https://doi.org/10.1016/j.ecoser.2016.11.011

Sushinsky, J.R., Rhodes, J.R., Possingham, H.P., Gill, T.K., Fuller, R.A., 2013. How should we grow cities to minimize their biodiversity impacts? Global Change Biology 19, 401–410. https://doi.org/10.1111/gcb.12055

Taguchi, V.J., Weiss, P.T., Gulliver, J.S., Klein, M.R., Hozalski, R.M., Baker, L.A., Finlay, J.C., Keeler, B.L., Nieber, J.L., 2020. It is not easy being green: Recognizing unintended consequences of green stormwater infrastructure. Water (Switzerland) 12, 522. https://doi.org/10.3390/w12020522

Talal, M.L., Santelmann, M. V., 2019. Plant community composition and biodiversity patterns in urban parks of Portland, Oregon. Frontiers in Ecology and Evolution 7. https://doi.org/10.3389/fevo.2019.00201

Tam, K.C., Bonebrake, T.C., 2016. Butterfly diversity, habitat and vegetation usage in Hong Kong urban parks. Urban Ecosystems 19, 721–733. https://doi.org/10.1007/s11252-015-0484-2

Tashiro, A., Nakaya, T., Nagata, S., Aida, J., 2021. Types of coastlines and the evacuees’ mental health: A repeated cross-sectional study in Northeast Japan. Environmental Research 196, 110372. https://doi.org/10.1016/j.envres.2020.110372

Thiagarajan, M., Newman, G., Van Zandt, S., 2018. The projected impact of a neighborhood-scaled green-infrastructure retrofit. Sustainability (Switzerland) 10, 3665. https://doi.org/10.3390/su10103665

Threlfall, C.G., Ossola, A., Hahs, A.K., Williams, N.S.G., Wilson, L., Livesley, S.J., 2016. Variation in vegetation structure and composition across urban green space types. Frontiers in Ecology and Evolution 4. https://doi.org/10.3389/fevo.2016.00066

Tiwari, A., Kumar, P., 2020. Integrated dispersion-deposition modelling for air pollutant reduction via green infrastructure at an urban scale. Science of the Total Environment 723, 138078. https://doi.org/10.1016/j.scitotenv.2020.138078

Tomson, M., Kumar, P., Barwise, Y., Perez, P., Forehead, H., French, K., Morawska, L., Watts, J.F., 2021. Green infrastructure for air quality improvement in street canyons. Environment International 146, 106288. https://doi.org/10.1016/j.envint.2020.106288

Tonietto, R., Fant, J., Ascher, J., Ellis, K., Larkin, D., 2011. A comparison of bee communities of Chicago green roofs, parks and prairies. Landscape and Urban Planning 103, 102–108. https://doi.org/10.1016/j.landurbplan.2011.07.004

Tryjanowski, P., Morelli, F., Mikula, P., Krištín, A., Indykiewicz, P., Grzywaczewski, G., Kronenberg, J., Jerzak, L., 2017. Bird diversity in urban green space: A large-scale analysis of differences between parks and cemeteries in Central Europe. Urban Forestry and Urban Greening 27, 264–271. https://doi.org/10.1016/j.ufug.2017.08.014

Tsai, W.L., Davis, A.J.S., Jackson, L.E., 2019. Associations between types of greenery along neighborhood roads and weight status in different climates. Urban Forestry and Urban Greening 41, 104–117. https://doi.org/10.1016/j.ufug.2019.03.011

Tsai, W.L., McHale, M.R., Jennings, V., Marquet, O., Hipp, J.A., Leung, Y.F., Floyd, M.F., 2018. Relationships between characteristics of urban green land cover and mental health in U.S. metropolitan areas. International Journal of Environmental Research and Public Health 15, 340. https://doi.org/10.3390/ijerph15020340

Tzortzakaki, O., Kati, V., Kassara, C., Tietze, D.T., Giokas, S., 2018. Seasonal patterns of urban bird diversity in a Mediterranean coastal city: the positive role of open green spaces. Urban Ecosystems 21, 27–39. https://doi.org/10.1007/s11252-017-0695-9

Van Mechelen, C., Van Meerbeek, K., Dutoit, T., Hermy, M., 2015. Functional diversity as a framework for novel ecosystem design: The example of extensive green roofs. Landscape and Urban Planning 136, 165–173. https://doi.org/10.1016/j.landurbplan.2014.11.022

Wallner, P., Kundi, M., Arnberger, A., Eder, R., Allex, B., Weitensfelder, L., Hutter, H.P., 2018. Reloading pupils’ batteries: Impact of green spaces on cognition and wellbeing. International Journal of Environmental Research and Public Health 15, 1205. https://doi.org/10.3390/ijerph15061205

Wood, L., Hooper, P., Foster, S., Bull, F., 2017. Public green spaces and positive mental health – investigating the relationship between access, quantity and types of parks and mental wellbeing. Health and Place 48, 63–71. https://doi.org/10.1016/j.healthplace.2017.09.002

Wu, L., Kim, S.K., 2021. Health outcomes of urban green space in China: Evidence from Beijing. Sustainable Cities and Society 65, 102604. https://doi.org/10.1016/j.scs.2020.102604

Wurth, A.M., Ellington, E.H., Gehrt, S.D., 2020. Golf Courses as Potential Habitat for Urban Coyotes. Wildlife Society Bulletin 44, 333–341. https://doi.org/10.1002/wsb.1081

Xie, C., 2018. Tree Diversity in Urban Parks of Dublin, Ireland. Fresenius Environmental Bulletin 27, 8695–8708.

Xie, S., Ouyang, Z., Gong, C., Meng, N., Lu, F., 2021. Seasonal fluctuations of urban birds and their responses to immigration: An example from Macau, China. Urban Forestry and Urban Greening 59, 126936. https://doi.org/10.1016/j.ufug.2020.126936

Yang, G., Xu, J., Wang, Y., Wang, X., Pei, E., Yuan, X., Li, H., Ding, Y., Wang, Z., 2015. Evaluation of microhabitats for wild birds in a Shanghai urban area park. Urban Forestry and Urban Greening 14, 246–254. https://doi.org/10.1016/j.ufug.2015.02.005

Yang, J., Sun, J., Ge, Q., Li, X., 2017. Assessing the impacts of urbanization-associated green space on urban land surface temperature: A case study of Dalian, China. Urban Forestry and Urban Greening 22, 1–10. https://doi.org/10.1016/j.ufug.2017.01.002

Yang, L., Zhang, L., Li, Y., Wu, S., 2015. Water-related ecosystem services provided by urban green space: A case study in yixing city (china). Landscape and Urban Planning 136, 40–51. https://doi.org/10.1016/j.landurbplan.2014.11.016

Zhang, D., Zheng, H., He, X., Ren, Z., Zhai, C., Yu, X., Mao, Z., Wang, P., 2016. Effects of forest type and urbanization on species composition and diversity of urban forest in Changchun, Northeast China. Urban Ecosystems 19, 455–473. https://doi.org/10.1007/s11252-015-0473-5

Zhang, X., Estoque, R.C., Murayama, Y., 2017. An urban heat island study in Nanchang City, China based on land surface temperature and social-ecological variables. Sustainable Cities and Society 32, 557–568. https://doi.org/10.1016/j.scs.2017.05.005

Zhu, Z.X., Pei, H.Q., Schamp, B.S., Qiu, J.X., Cai, G.Y., Cheng, X.L., Wang, H.F., 2019. Land cover and plant diversity in tropical coastal urban Haikou, China. Urban Forestry and Urban Greening 44, 126395. https://doi.org/10.1016/j.ufug.2019.126395

Zölch, T., Henze, L., Keilholz, P., Pauleit, S., 2017. Regulating urban surface runoff through nature-based solutions – An assessment at the micro-scale. Environmental Research 157, 135–144. https://doi.org/10.1016/j.envres.2017.05.023

Zölch, T., Maderspacher, J., Wamsler, C., Pauleit, S., 2016. Using green infrastructure for urban climate-proofing: An evaluation of heat mitigation measures at the micro-scale. Urban Forestry and Urban Greening 20, 305–316. https://doi.org/10.1016/j.ufug.2016.09.011
