## Appendix 5 for "CUGIC: The Consolidated Urban Green Infrastructure Classification for assessing ecosystem services and biodiversity"

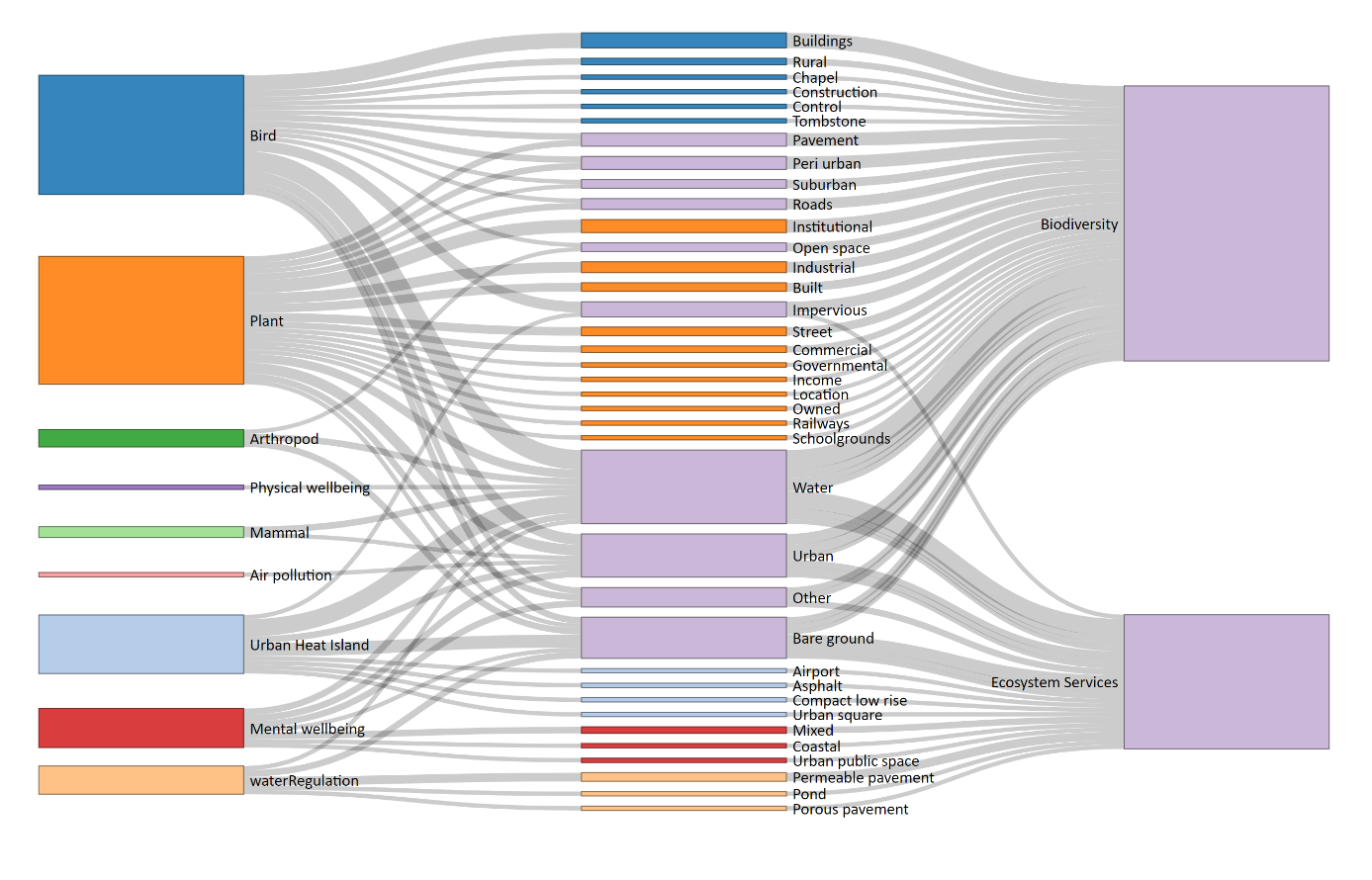

**Appendix 3A. Sankey diagram showing the links between services or taxon, grey and blue infrastructure class names, and discipline.** Nodes in the middle are ordered with the least frequently used grey and blue infrastructure classes on the edges, while most frequently used classes are in the center. The thickness of the nodes is based on frequency.

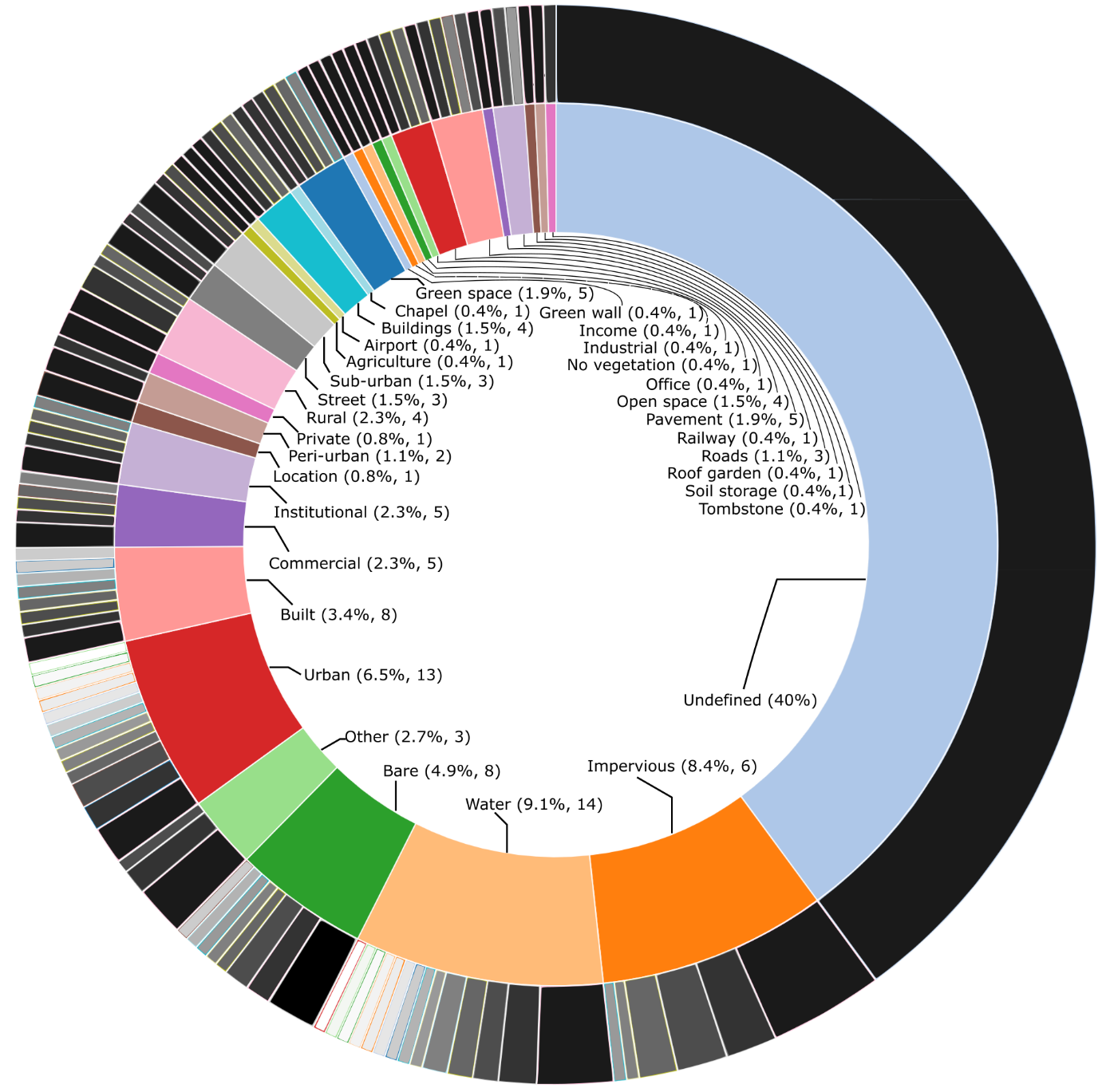

**Appendix 3B. Sunburst diagram of the grey and blue infrastructure themes and types.** The inner layer contains themes, and the outer layer types, portraying the number of grey and blue infrastructure types in one theme.

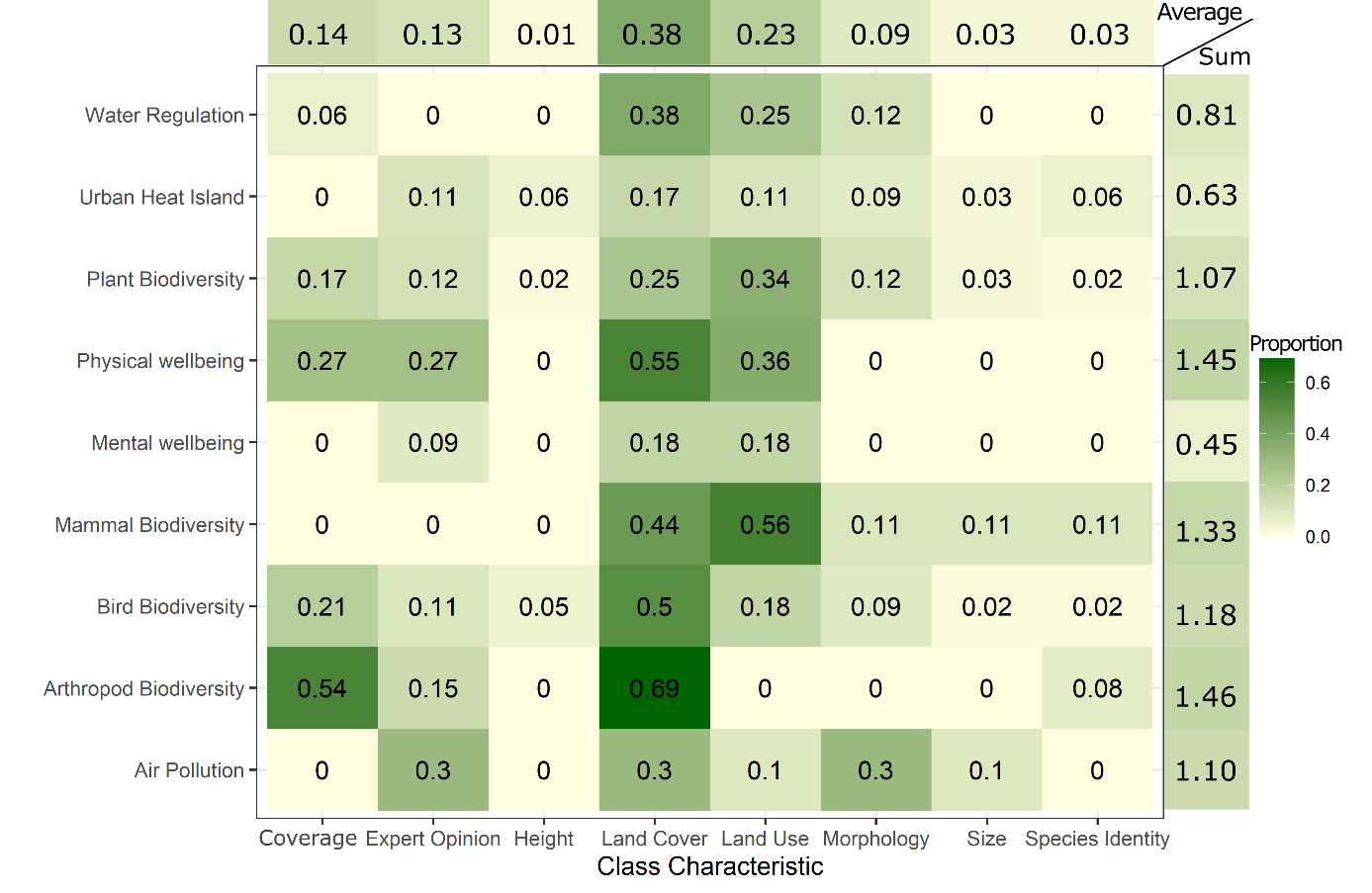

**Appendix 3C. Heat map with proportional characteristics of grey and blue infrastructure classes by ES and taxa.** A higher value indicates that proportionally more classes have this characteristic. Averaged and summed values are shown at end of the matrix’s columns and rows. Average values indicate use of one characteristics per ES or taxon (0-1) and summed scores indicate the average use of all characteristics in a ES or taxon (0-8).
