## Appendix 6 for "CUGIC: The Consolidated Urban Green Infrastructure Classification for assessing ecosystem services and biodiversity"

***Appendix 5. Details on binary scoring of class characteristics***

Eight class characteristics were scored binary based on presence or absence. Land use or land cover were scored 1 if they were explicitly mentioned. Size, height, or coverage were scored 1 if the definition explicitly mentioned either square meters, meters of height or percentage of coverage, respectively. Morphology was scored 1 if the form of the GI, or its place in respect to other structures, was explicitly stated. Species identity was scored 1 when it specified a particular species, or if it specified a particular trait of species (e.g. evergreen, deciduous, etc.). We chose to not score grass, forb, tree and other general names of vegetation as a 1 for species identity due to ambiguity of their definition across services or taxa (i.e. grass could imply species or vegetation below 0.5m). Finally, expert opinion was scored 1 when a class was based on photographs, locations, expert opinion, or machine learning algorithms. This meant that a study defining natural as “unsettled lands that may occasionally include dwellings. 0-2% built; 0 buildings/ha; <1 human /ha” would score 1 on land-use and coverage as it provides explicit information on the use of land and the percentage coverage by built-up area, while a score of 0 was given for the other elements as they are not explicitly defined.
