## Appendix 7 for "CUGIC: The Consolidated Urban Green Infrastructure Classification for assessing ecosystem services and biodiversity"

**Appendix 7. Table containing the changes from the Green Infrastructure (GI) themes to CUGIC categories.** Shown in the left two columns are the original GI themes as consolidated through the data acquired from the literature. The right two columns shows the resulting CUGIC class, and motivation for any changes made.

| **Theme** | **Description** | **CUGIC Layer** | **CUGIC class** | **Motivation for changes** |
| --- | --- | --- | --- | --- |
| Natural | Land maintained in its natural or native state, if any land-use its aim to conserve biodiversity. Little to no management of the vegetation. Recreation is limited to passive use, such as benches for sitting after walking. This can involve, but is not limited to; woodlands, scrub, grasslands, coastal regions, etc. | LULC | Natural | Removed the aim to conserve biodiversity, as it is hard to quantify or decide to what extent the intent is. Mainly we now define it as “Land maintained in its natural or native state. Little to no management of the vegetation. Recreation is limited to passive use, such as benches for sitting after walking. This can involve, but is not limited to; woodlands, scrub, grasslands, coastal regions, etc.” Vegetation difference are further detailed in the vegetation layer of the CUGIC |
| Lot | land without evident use, leading to unmanaged vegetation growth. | LULC | Remnant vegetation | All three theme converge on having no evident management leading to remnant vegetation. Without having an easy distinction between them, we argue that it is better to combine them.  Combined as they are often hard to differentiate. |
| Spontaneous | Vegetation without evident management, usually grown spontaneous | LULC | Remnant vegetation |  |
| Remnant | Vegetation without evident management | LULC | Remnant vegetation |  |
| Park | Green spaces designated for a wide variety of passive and active recreation. Are usually designed and intensely managed. | LULC | Park | No changes made |
| Agriculture | Land used in the broadest sense of agriculture. This includes cash-crops, horticulture, pastoral uses, orchard, etc. Usually fairly large in size in comparison to other land-uses. | LULC | Agriculture | No changes made |
| Botanical Garden | Garden used for scientific purposes | LULC | Botanical Garden | No changes made |
| Cemetery | Land used as cemetery involves low management of the vegetation. Always contain trees. Land-use is unaltered for a long periods of time. Can be small or large. | LULC | Cemetery | No changes made |
| Golf course | Land designed for golf uses. This consists of a grassy environment with at their periphery natural (unmanaged) landscape (usually forest). | LULC | Golf course | No changes made |
| Bioswale | Street scale linear vegetated mulches with a gentle slope. Usually have an engineered soil. Placed next to impervious environment such as roads or parking spaces. | LULC | Bioswale | No changes made |
| Wetlands | Engineered systems that regulates water next to permanent water bodies | LULC | Wetland | No changes made |
| Detention basin | No clear definition among definitions | LULC | Detention basin | After definition search, we provided the next definition: “land excavated next to flooding water bodies” |
| Infiltration basin | No clear definition among definitions | LULC | Infiltration basin | After definition search, we provided the next definition: “Constructed apparatus to regulate water” |
| Bioretention basin | No clear definition among definitions | LULC | Rain garden | Bioretention basin is a synonym for rain garden. |
| Rain garden | Trees or vegetation next to impermeable surface with permeable earth | LULC | Rain garden | No changes made |
| Residential | Vegetation in private yards next to or close to residential housing. | LULC | Residential green & garden | No changes made |
| Garden | Generally decorative purposes, and thus often contains flowerbeds or flowering species. | LULC | Residential green & garden | No changes made |
| Green corridor | Strips of vegetation. Usually involves grass and scattered trees, can involve shrubs. In regards to other infrastructure, they are usually placed next to canals, roads, cycling path, etc. | LULC | Green buffer zones | Changed name, kept content the same. |
| Green roof | Vegetation that partially or fully covers a roof | LULC | Green roof | No changes made |
| Vertical green | Vegetation along a vertical surface. These can involve climbing plants, green walls, green facade, etc. | LULC | Green wall | No changes made |
| Height | Any height created by vegetation layers. Involves a broad spectrum of height requirements. | Vegetation | Vegetation categories | We have chosen to use the two most commonly used height thresholds being 1m and 5m. Most studies, aim to quantify lower, mid, and high vegetation in this way. That broadly reflects forests, shrubs, and grass/herb vegetation. |
| Layered | Contains either one or multiple layers of vegetation | Vegetation | Vegetation categories | We have chosen to include vegetation being either layered or not. This can inform whether there are solely trees or shrubs, or also lower levels of vegetations involved. |
| (Tree) Canopy | Any form of canopy created by either trees or shrubs. Involves a broad spectrum of coverage % | Vegetation | Vegetation categories | We have chosen to use the three most commonly used vegetation coverage threshold, being 10%, 50%, and 70%. |
| Wooded | Any land with substantial and continuous cover by woody vegetation. Can be a mix of coniferous, deciduous and evergreen species. | Vegetation | Vegetation categories | We argue that the often used evergreen forest or deciduous forest informs one about potential temporally important effects. Therefore we have decided to additionally include on top of the regular vegetation categories for the forest categories (i.e. vegetation above 5m). |
| Shrub | Shrub species | Vegetation | Vegetation categories | These themes largely rely on species identity. This is hard to infer from remote sensing data. Thus we have chosen to include them through the vegetation categories create by the themes above, being height & vegetational coverage. From this we infer by height which kind of vegetation it is, and how dense it is. |
| Herb | Grass or herbaceous species | Vegetation | Vegetation categories |  |
| Grassland | Land covered primarily with grasses, but also includes other low level vegetation (<0.5m). Can sparsely have some trees. | Vegetation | Vegetation categories |  |
| Green open space | Land uses that are comprised mainly of low level vegetation, and without any substantial tree cover. This involves for example, parks, gardens, grassland, cemeteries and recreational grounds. The main purpose of the green space is for human land uses. Usually administered by the municipality and public. | Excluded | Excluded | Green open space are, in our results, used as a way to define large green spaces. This could almost be anything, and aggregates many of the other themes to a coarser scale. We have opted to exclude it, as it does not provide additional detail to assessing biodiversity or ES in future studies. |
| Courtyard | Courtyard with grass and or a tree. Used solely in two studies. Not used further. | Excluded | Excluded | Used in solely two VR studies. |
